# Ethanol Drives Evolution of Hsp90-Dependent Robustness by Redundancy in Yeast Domestication

**DOI:** 10.1101/2023.07.21.547572

**Authors:** Dipak Patel, Hatim Amiji, William Shropshire, Natalia Condic, Nejla Ozirmak Lermi, Youssef Sabha, Beryl John, Blake Hanson, Georgios Ioannis Karras

## Abstract

Protein folding promotes and constrains adaptive evolution. We uncover this surprising duality in the role the protein-folding chaperone Hsp90 plays in mediating the interplay between proteome and the genome which acts to maintain the integrity of yeast metabolism in the face of proteotoxic stressors in anthropic niches. Of great industrial relevance, ethanol concentrations generated by fermentation in the making of beer and bread disrupt critical Hsp90-dependent nodes of metabolism and exert strong selective pressure for increased copy number of key genes encoding components of these nodes, yielding the classical genetic signatures of beer and bread domestication. This work establishes a mechanism of adaptive canalization in an ecology of major economic significance and highlights Hsp90-contingent variation as an important source of phantom heritability in complex traits.

## Introduction

Biological systems are remarkably robust to internal and external fluctuations but the origins of such widespread robustness have remained enigmatic. Classical experiments conducted by C.H. Waddington revealed that the robustness of developmental traits to environmental variations is in part genetically determined and can be selected, a phenomenon he coined “canalization” (Waddington 1942, Scharloo 1991). Since Waddington’s observation, canalization has gone on to captivate the imagination of many and spark a revolution in biology with far-reaching implications extending from human health to ecosystem sustainability and food security to biotechnology (Alon 2003, Abson, Fraser et al. 2013, Gong, Nielsen et al. 2017, Gibson and Lacek 2020, Crespi, Burnap et al. 2022). However, despite the long history of the concept, the proximal (molecular) and evolutionary mechanisms driving robustness and canalization remain obscure and their practical applications unexploited (Gibson and Wagner 2000, de Visser, Hermisson et al. 2003, Stelling, Sauer et al. 2004, Flatt 2005, Hallgrimsson, Green et al. 2019). Without deep knowledge of these mechanisms, evolution will remain difficult to explain or harness for human benefit in medicine and industry alike.

Many mechanisms have been shown to promote robustness, sometimes adaptively (Montville, Froissart et al. 2005, Sanjuan, Cuevas et al. 2007, McBride, Ogbunugafor et al. 2008, Goldhill, Lee et al. 2014), and these mechanisms can differ depending on the source of perturbation (e.g. robustness against mutations) (Scharloo 1991, Wagner, Booth et al. 1997, Kitano 2004, Felix and Barkoulas 2015, Payne and Wagner 2019). Diverse unrelated genes, especially those encoding proteins at the center of intracellular network hubs, can “buffer” both genetic and environmental variation as well as biological noise, thereby contributing to canalization (de Visser, Hermisson et al. 2003, Wagner 2005). Perhaps the best studied genes underlying buffering mechanisms encode members of the heat shock protein 90 (Hsp90) family (Zabinsky, Mason et al. 2019). Hsp90 is an ancient and highly conserved ATP-dependent protein-folding chaperone that evolved to safeguard protein homeostasis within the cell by assisting in the folding and maturation of diverse protein substrates (“clients”). Unlike the other major general chaperones in the cell, Hsp90 is constitutively expressed in excess over cellular requirements for growth and survival (Borkovich, Farrelly et al. 1989, Bhattacharya, Maiti et al. 2022). Excess Hsp90 provides a protein-folding buffer to the cell that supports many cellular pathways and has been shown to influence the relationship between genotype and phenotype (Rutherford and Lindquist 1998, Fares, Ruiz-Gonzalez et al. 2002, Gorre, Ellwood-Yen et al. 2002, Nimmanapalli, O’Bryan et al. 2002, Queitsch, Sangster et al. 2002, Cowen and Lindquist 2005, Maisnier-Patin, Roth et al. 2005, Yeyati, Bancewicz et al. 2007, Bogumil and Dagan 2010, Jarosz and Lindquist 2010, Casanueva, Burga et al. 2012, Rohner, Jarosz et al. 2013, Aguilar-Rodriguez, Sabater-Munoz et al. 2016, Kadibalban, Bogumil et al. 2016, Karras, Yi et al. 2017, Iyengar and Wagner 2022, Iyengar and Wagner 2022). Importantly, the Hsp90 buffering capacity of the cell is finite and can be exceeded by diverse proteotoxic challenges which lead to protein unfolding and the titration of Hsp90 away from its normal clients with consequent loss of their function (Zabinsky, Mason et al. 2019). This aspect of Hsp90 function allows the chaperone to couple the effects of genetic variation to diverse proteotoxic cues in the environment of the cell.

Several independent lines of evidence suggest that Hsp90-contingent genetic variation has adaptive value (Queitsch, Sangster et al. 2002, Cowen and Lindquist 2005, Whitesell and Lindquist 2005, Sangster, Salathia et al. 2008, Jarosz and Lindquist 2010, Rohner, Jarosz et al. 2013, Whitesell, Santagata et al. 2014, Karras, Yi et al. 2017). In enabling evolution, Hsp90 can act as a capacitor or a potentiator of genetic variation. As a capacitor, Hsp90 mitigates the effects of genetic variation thus enabling its accumulation within populations (Rutherford and Lindquist 1998, Queitsch, Sangster et al. 2002). Adverse exposures can reveal Hsp90-buffered mutations, thus exposing them to natural selection. As a potentiator, Hsp90 allows mutants to express new phenotypic effects (Whitesell, Mimnaugh et al. 1994, Cowen and Lindquist 2005). However, mechanisms of robustness, including the Hsp90 buffer, have been traditionally studied in model traits in the laboratory or *in silico* mostly in isolation and independently of the ecological forces shaping their evolution in nature (Wagner, Booth et al. 1997, Wilkins 1997, Keane, Toft et al. 2014, Felix and Barkoulas 2015, Naseeb, Ames et al. 2017, Hallgrimsson, Green et al. 2019, Crespi, Burnap et al. 2022) with few exceptions (Cowen and Lindquist 2005, Rohner, Jarosz et al. 2013). Hence, the ecological stressors that reveal Hsp90-buffered variation and the proximal mechanisms driving the evolution of robustness in nature remain elusive. It is also unknown how different mechanisms of robustness interact. Furthermore, the adaptive value of robustness is debated; that is whether robustness against mutations (genetic canalization) is the target of natural selection or evolves congruently by selection for robustness to environmental stress (environmental canalization) or as a pleiotropic byproduct of other adaptations or yet again as an unavoidable consequence of increased complexity (Wagner, Booth et al. 1997, Bergman and Siegal 2003, Harrison, Papp et al. 2007, Wagner and Zhang 2011, Keane, Toft et al. 2014, Felix and Barkoulas 2015, Naseeb, Ames et al. 2017, Crespi, Burnap et al. 2022).

To tackle our limited understanding of Hsp90-dependent robustness in evolution, we decided to begin by focusing on yeast metabolism. Metabolic traits have enabled budding yeast species to spread across diverse ecological niches and in some cases have been strongly selected in man-made niches (Warringer, Zorgo et al. 2011, Goddard and Greig 2015). Yeast as a model provides a rich resource of domesticated and wild strains to study evolution especially for metabolism which is a model system in the study of robustness and Hsp90 is known to support key nodes in the metabolic network of yeast such as Gal1, Mal63 (Bali, Zhang et al. 2003, Gopinath and Leu 2016). We employed Hsp90 inhibition as a proxy for proteotoxic environmental stress—and evaluated differences in the sensitivity of Hsp90-dependent metabolic traits across 698 yeast strains of diverse domesticated and wild origins. Our results demonstrate that industrially relevant metabolic traits in beer and bread yeasts have evolved robustness to stresses which exceed Hsp90 chaperone capacity. Gene duplications characteristic of yeast domestication in maltose-rich niches stabilize desirable industrial traits against Hsp90 stressors that are prevalent in the industrial niche. As a telling example, ethanol exerts selective pressure for gene duplications conducive to trait canalization, leading us to propose that industrial traits have undergone genetic and environmental canalization through selection for robustness against Hsp90 stress during domestication of yeast for beer and bread production.

## Results

### Robustness of metabolic traits to Hsp90 stress is rapidly evolving

To investigate the role of Hsp90 in the evolution of yeast metabolism we evaluated 11 metabolic traits across diverse strains from 10 species of the *Saccharomyces sensu stricto* clade, including 79 *S. cerevisiae* and 83 *S. paradoxus* isolates. To allow quantitative comparisons for each metabolic trait, we generated duplicate growth curves in high throughput using media containing a single carbohydrate as the sole source of carbon. This revealed substantial variation between different strains and growth conditions (Figures 1A and S1A). The area under the curve integrates multiple growth parameters and was therefore chosen as a proxy for relative growth (Figure S1A). Pooled relative growth values across the examined strains and conditions examined were mostly bimodal in distribution, with a small number of intermediate values observed (Figure S1B). Normalization of relative growth estimates for each condition over glucose provided a proxy for relative metabolic efficiency (Figure S1C).

**Figure 1.**
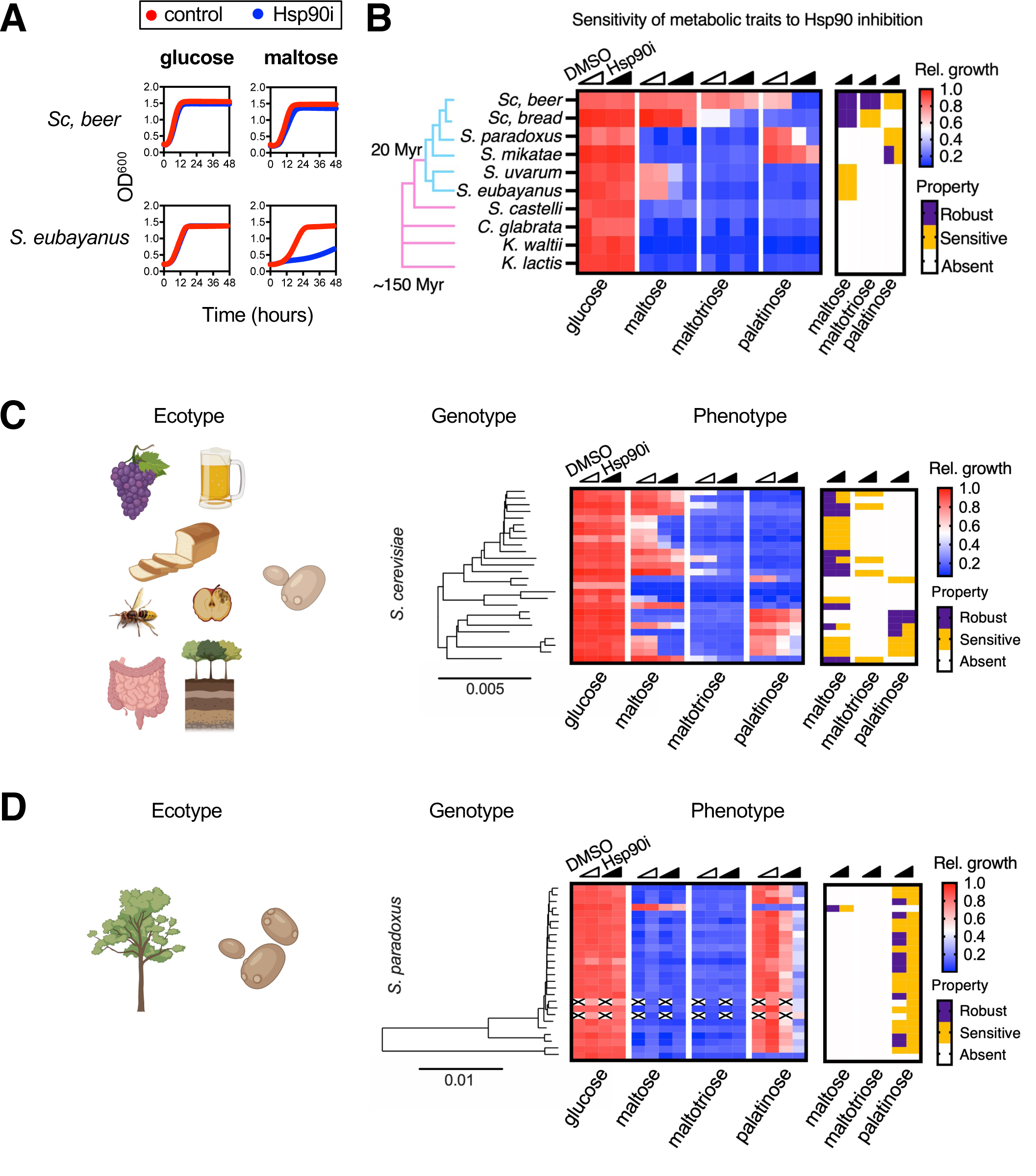
Ecology shapes trait robustness to Hsp90 stress. (A) Representative growth curves for a *S. pastorianus* lager beer strain and a *S. eubayanus* wild isolate growing in glucose and maltose with or without radicicol (10 μM). Standard deviations are smaller than the size of the symbol. (B-D) Heat maps of relative growth and trait robustness to Hsp90 impairment derived from growth curves in rich media containing the indicated carbohydrate as the sole source of carbon are shown for (B) 10 *Saccharomyces sensu stricto* strains, (C) 26 *S. cerevisiae* MAT alpha isolates, and (D) 26 MAT alpha *S. paradoxus* isolates. Filled triangles indicate samples exposed to increasing concentrations of radicicol. Empty triangles indicate samples treated with vehicle control (DMSO). Robustness to Hsp90 inhibition (radicicol, 4 μM *vs.* 10 μM) was color-coded based on a threshold of −0.1 (orange, robustness low, ≤ −0.1; violet, robustness high, > −0.1; white, no growth or not determined). Depictions of the ecological niches from which the species have been isolated (ecotypes). Dendrograms of yeast genomes, with estimated time of divergence to the whole-genome duplication event indicated (Myr, million years). Values represent averages of 2 – 3 independent experiments. Pictograms were created with BioRender (https://biorender.com/).

The efficiency of metabolic traits varied drastically across *Saccharomyces sensu stricto* yeasts (Figures 1B and S1D). We also observed marked variation in metabolism across diverse *S. cerevisiae* ecotypes, especially for galactose, and maltose metabolism (Figures 1C and S1D). Strikingly, *S. paradoxus* isolates predominantly isolated from oak trees utilized carbohydrates much more uniformly, as compared to *S. cerevisiae* strains (F-test *p* << 0.005; Figures 1D and S1D). *S. paradoxus* strains shared a strong preference for turanose, palatinose, methyl-α-D-glucopyranoside (methyl-αG) and melezitose, which are sugars reported to be prevalent in sap and other related ecological niches which this species inhabits (Behera and Balaji 2021). These results were highly reproducible between technical replicates (two-tailed Pearson correlation, R^2^ > 0.99, *p* < 0.0001), individual colonies (two-tailed Pearson correlation, R^2^ > 0.99, *p* < 0.0001) and independent experiments (two-tailed Pearson correlation, R^2^ > 0.98, *p* < 0.0001; Figure S1E), and agree with previous findings supporting the idea that variation in metabolic traits is largely defined by population history and ecology in yeast (Warringer, Zorgo et al. 2011). Hence, we have identified a panel of rapidly evolving and ecologically relevant traits with which to probe Hsp90’s role in the evolution of metabolism.

Next, we evaluated the effects of Hsp90 impairment on each one of the above-mentioned strain/conditions. To compromise Hsp90 function, we supplemented each culture with the naturally occurring resorcinylic Hsp90 inhibitor, radicicol, which induced low or moderate Hsp90 stress at concentrations of 4 μM and 10 μM concentrations, respectively (Jarosz and Lindquist 2010). At both concentrations, radicicol perturbed diverse metabolic traits without affecting growth in the presence of glucose (Figure S1F). These effects were highly reproducible (two-tailed Pearson correlation, technical replicates: R^2^ > 0.99, *p* < 0.0001, individual colonies: R^2^ > 0.98, *p* < 0.0001, replicate experiments: R^2^ > 0.95, *p* < 0.0001; Figure S1E). In addition, the results were unaffected by mating type or the number of generations over which the cells had been in continuous culture (two-tailed Pearson correlation, R^2^ > 0.96, *p* < 0.0001; Figure S1G). Structurally diverse small-molecule Hsp90 inhibitors reproduced the effects of radicicol (two-tailed Pearson correlation, *p* < 0.0001; Figures S1H and S1I), corroborating our finding that metabolic traits are sensitive to Hsp90 inhibition. Furthermore, expression of hypomorphic Hsp82 mutants, A587, G313S, T101I, as sole source of Hsp90 protein (Nathan and Lindquist 1995) impaired palatinose metabolism (Figure S1J). The Hsp82-A587T mutant exerted strong effects on palatinose metabolism without significantly affecting glucose utilization under basal conditions yet showed increased sensitivity to radicicol (Figure S1J). Moreover, overexpression of the constitutive Hsp90 isoform, Hsc82, allowed robust palatinose metabolism even upon Hsp90 inhibition (Figure S1J). We conclude that radicicol exposure perturbs metabolic traits by causing Hsp90 impairment.

Examining the dose-response relationship for radicicol and palatinose metabolism in a laboratory strain revealed differences in growth < 5% (Hedge’s *g* effect size > 1). The extracted estimates of metabolic efficiency and robustness to various strengths of Hsp90 inhibition revealed greater effect on trait mean than trait variance and were all normally distributed (Figure S1K). To ensure physiological relevance, we applied a stringent significance threshold for metabolic robustness against Hsp90 inhibition (radicicol vs. vehicle control, Log_10_ = −0.1, effect size > 4; Figure S1L). Applying this analysis to each strain-trait pair with metabolic efficiency greater than 25% (“positive”; Figure S1C) revealed that 35% of all strain-trait interactions were sensitive to Hsp90 inhibition (Figure S1M); approximately 3 out of 4 strains expressed at least one Hsp90-dependent metabolic trait. Robustness estimates were highly reproducible between replicates (Figure S1N). These experiments identified diverse metabolic traits that were sensitive to Hsp90 inhibition, including maltose, maltotriose, and palatinose metabolism (Figures S1F and S1O), suggesting a broader role for Hsp90 in the evolution of peripheral carbon metabolism in yeast.

Importantly, the sensitivity of metabolic traits to Hsp90 stress−used henceforth to denote sub-toxic impairment of Hsp90 function−varied drastically between closely related yeast species and even between intra-species isolates (Figure S1O). In contrast, the sensitivity of palatinose metabolism to Hsp90 impairment was strikingly uniform across *S. paradoxus* strains (Figures 1C and S1O). The observed variance in trait robustness to Hsp90 impairment reflects ecological differences between the domesticated *S. cerevisiae* and the non-domesticated *S. paradoxus* yeast species.

### Trait robustness to Hsp90 stress as a new phenotypic signature of beer and bread yeasts

Domestication is a recent event in the evolutionary history of budding yeasts that is continuing in the modern day with the extensive industrialization of fermentation processes (Steensels, Gallone et al. 2019, Lahue, Madden et al. 2020). To determine how yeast domestication has influenced the robustness of metabolic traits to sub-toxic Hsp90 stress, we evaluated the sensitivity of two industrially relevant metabolic traits, maltose and maltotriose metabolism, to Hsp90 inhibition across 534 natural isolates of ecologically diverse origins (Figure 2A). This library included 321 domesticated strains (i.e., used in beer brewing, wine making, baking and other food-related fermentative processes), 32 strains of unknown origin and a reference of 181 non-domesticated wild isolates (i.e., clinical, soil, wasp gut-derived). Additionally, we examined the ability of these strains to utilize palatinose, a genetically distinct trait yet one with a genetic composition comparable to maltose and maltotriose metabolism (Brown, Murray et al. 2010). Since palatinose metabolism is an Hsp90-dependent and ecologically divergent metabolic trait, palatinose results served as an independent non-domesticated control (Figures 1A-C and S1L-O).

**Figure 2.**
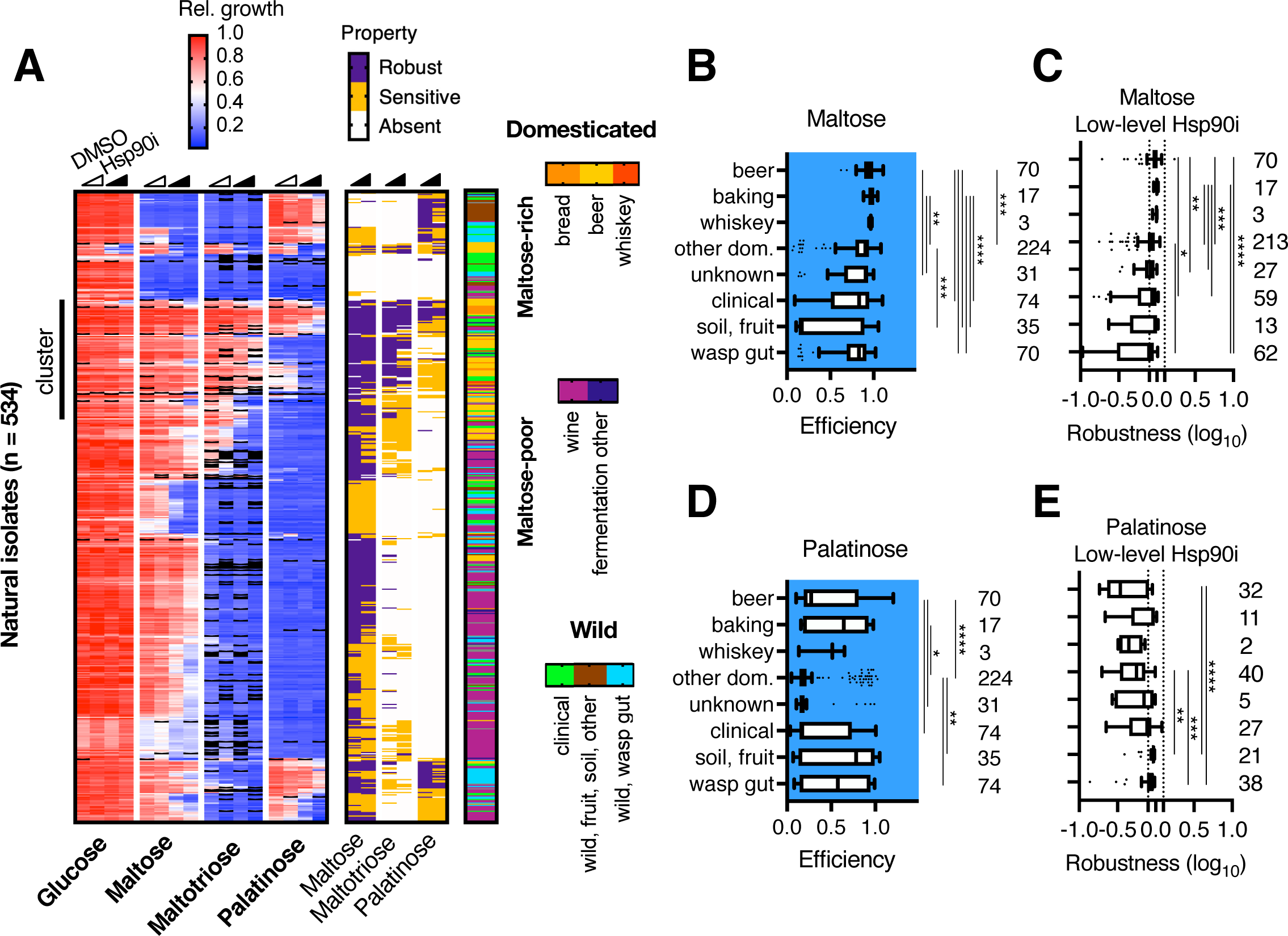
Selection for trait robustness to Hsp90 stress during yeast domestication. (A) Heat map of relative growth and robustness estimates for 534 natural *Saccharomyces* isolates determined for the indicated metabolic traits under basal conditions (empty triangles) and under low and moderate radicicol concentrations (Hsp90i, filled triangles). Strain origin is indicated on the right. (B-E) Efficiency of (B) maltose and (D) palatinose metabolism and robustness of these traits to low-level Hsp90 inhibition (4 μM; C and E). Values represent averages of 2 – 3 independent experiments. Whiskers based on Tukey method. Significance between distributions was determined using the Kruskal-Wallis test with Dunn’s multiple comparisons.

We generated 33,553 growth curves examining maltose, maltotriose and palatinose metabolism, across the 534 natural yeast isolates, under conditions of basal *versus* low- and moderate-level Hsp90 inhibition. Experiments were conducted in technical duplicate and in independent triplicate experiments (Figure 2A). The distributions of efficiencies and robustness estimates within this larger dataset were comparable to the results we obtained for wild-derived strains (compare Figure S2B to S1C, and Figure S2C to S1M). These results were highly reproducible between technical and biological replicates (Figure S2D) as well as with a structurally distinct Hsp90 inhibitor (NVP-HSP990, 25 μM, two-tailed Pearson correlation, R^2^ = 0.9053, *p* < 0.0001; Figure S2E). Overall, of the ∼ 55% natural isolates that could metabolize maltose, maltotriose or palatinose, > 83% were sensitive to Hsp90 inhibition for at least one metabolic trait, and > 50% of the examined strain-trait interactions were perturbed by Hsp90 inhibition (Figures 2A and S2C). These findings strongly support an evolutionary role for Hsp90 as potentiator of yeast metabolic traits.

Next, we examined associations between yeast domestication and the efficiency or robustness of metabolic traits to Hsp90 impairment. In accordance with an adaptive role for maltose and maltotriose metabolism in yeast domestication (Gallone, Steensels et al. 2016), domesticated strains metabolized these sugars with greater efficiency as compared to wild isolates (left panel; Figure S2F). Notably, the robustness of maltose but not maltotriose metabolism was associated with domestication (right panel; Figure S2F). However, there was weak correlation between the two traits regarding both efficiency and robustness to Hsp90 impairment (two-tailed Pearson correlation, R^2^ = 0.2426 and 0.1945, respectively, *p* < 0.0001; Figure 2A). Yet, the ability to metabolize maltotriose, not maltose, is the hallmark trait of domesticated beer yeasts (Gallone, Steensels et al. 2016). This is surprising because maltotriose not only coexists with maltose across diverse niches but is also less abundant than maltose in these environments.

Hierarchical clustering revealed a positive association between the basal efficiency of maltotriose metabolism and the robustness of maltose metabolism to Hsp90 impairment (Figures 2A, S2G and S2H). Consistent with this relationship, basal maltotriose metabolism correlated much better with maltose metabolism under Hsp90 inhibition (two-tailed Pearson correlation, R^2^ = 0.361, *p* < 0.0001) as compared to basal conditions (two-tailed Pearson correlation, R^2^ = 0.2426, *p* < 0.0001). We identified a cluster of maltotriose-utilizing strains that efficiently metabolized maltose even under Hsp90 inhibitory conditions (Figure 2A). Diverse strains, including beer yeasts, domesticated in maltose-rich niches (i.e., dough, wort) were enriched within this cluster (hypergeometric distribution tests for each strain class using sliding thresholds for robustness −0.079 – −0.699, *p* < 0.05; Figure 2A). Notably, bread and whiskey strains metabolized both maltose and maltotriose with comparable or superior efficiency compared to beer yeasts even under conditions of Hsp90 stress (Figures 2B and S2I). Indeed, both the basal efficiency of maltose and maltotriose metabolism and the robustness of both traits to Hsp90 impairment were greatly increased across domesticated strains from maltose-rich niches (i.e., beer, bread, whiskey), as compared to domesticated strains from maltose-poor environments (i.e., wine, sake, cider) (Figures 2B-C, S2I and S2J). We also observed substantial differences in the robustness of maltotriose, but not maltose, metabolism between ale and lager beer strains (Figure S2K). Differences in robustness over the entire dataset were largely independent of variation in basal metabolic efficiency (Figure S2L). The effects of Hsp90 inhibition were reproducible at different maltose concentrations (Figure S2M), including those in which the sugar is found in wort (Patel and Ingledew 1973), as well as in brewer’s wort instead of standard media (R^2^ = 0.801, *p* < 0.0001; Figure S2N).

In contrast to our results with maltotriose, we found no significant association between the robustness of maltose metabolism to Hsp90 inhibition and the basal efficiency of palatinose utilization (Figure S2F). Palatinose metabolism was highly sensitive to Hsp90 impairment across domesticated and wild isolates and comparable in basal efficiency and robustness to Hsp90 stress across beer, bread and other domesticated and wild strains (Figures 2D-E, S2H and S2J). A notable exception involved certain wild strains that were able to utilize palatinose in the face of Hsp90 stress (soil, fruit, wasp gut; Figure 2E). Hence, unlike maltose and maltotriose metabolism, the robustness of palatinose metabolism, albeit rapidly evolving, is not associated with beer and bread domestication. We conclude this evolved robustness or “canalization” of industrially relevant traits against compromise of Hsp90 function by factors in the environment is a novel phenotypic signature of beer and bread yeast domestication.

### Overlap between genetic signatures of yeast domestication and robustness to Hsp90 stress

Next, we asked whether the increased robustness of maltose metabolism to Hsp90 stress across beer and bread yeast isolates is genetically determined. To this end, we performed 11 crosses between diverse wild-derived *S. cerevisiae* haploid strains and the BY4741 laboratory strain which does not metabolize maltose, 7 of which did so in a manner robust to Hsp90 impairment. Inheritance of this trait followed a 2:2 Mendelian pattern in at least 7 of the 11 unique crosses, without bias towards trait robustness, with the parental pattern of robustness to Hsp90 stress being inherited in 125 of 168 dissected tetrads (74.4%,; Figure S3A). These findings suggest that the efficiency and robustness of metabolic traits are quantitative traits largely determined by a small number of metabolic genes. Because the robustness of maltose metabolism against Hsp90 impairment varied discordantly from the robustness of other traits across strains co-expressing 2 or more metabolic traits (Figure S3B), we concluded that trait robustness to Hsp90 stress is an intrinsic property of how the trait is supported in each strain, rather than an indirect manifestation of altered permeability of cells to Hsp90 inhibitors or differences in Hsp90 buffering capacity between strains.

To probe the genetic architecture of metabolic trait robustness to Hsp90 stress across industrial strains we performed whole-genome sequencing of 64 natural strains, parsed into 2 groups based on their sensitivity of maltose utilization to Hsp90 inhibition (radicicol, 10 μM; 32 robust and 32 sensitive). These strains include 18 bread strains, 13 beer strains, 8 distillery strains, and 25 wild isolates which served as a reference for non-domestication. The above strains were chosen based on their ability to metabolize maltose with high efficiency (>70%) and because each is representative of the variance in basal efficiency and robustness to Hsp90 stress of metabolic traits within each cognate group of yeast strain origin (Figure S3C). We sequenced these strains at 242x median coverage (min: 133.2x, max: 386.7x) in their original ploidy. Coverage at sub-telomeric regions varied by 2-4-fold between robust and sensitive groups (Figure S3D). Duplicated regions were associated with carbohydrate metabolism and maltose utilization gene terms (Figure S3E). Specifically, *MAL31* and *MAL32* and other metabolic genes were significantly amplified across strains that efficiently metabolized maltose under conditions of Hsp90 stress (Figure S3F). These genes encode permeases (*MALT* family) and hydrolases (*MALS* family), respectively (*MAL* genes), which are critical for the utilization of maltose and maltotriose (Brown, Murray et al. 2010, Duval, Alves et al. 2010). We found similar associations for genes in close linkage with *MAL31* and *MAL32 (YBR298C-A* and *YBR300C)*, genes containing repetitive sub-telomeric sequences (*YHL050C*, *YHR218W* and *YRF1-7*) and transposable element genes (*YLR410W-A*, *YCL019W*, and *YDR034C-C*) (Figure S3F). An independent Copy Number Variation (CNV) analysis algorithm produced comparable results (Control-FREEC *vs.* CONVICT, R^2^ = 0.9828, *p* < 0.0001; Figure S3G). These findings suggest that the robustness of maltose metabolism to Hsp90 stress is linked to duplications in the *MAL3* cluster and to a lesser extent in the *MAL1* cluster. These sub-telomeric *MAL* clusters contain *MALT* and *MALS* genes reported to comprise a genomic signature of domestication of beer and bread production (Michels and Needleman 1984, Day, Higgins et al. 2002, Dunn, Richter et al. 2012, Gallone, Steensels et al. 2016, Goncalves, Pontes et al. 2016, Duan, Han et al. 2018, Bigey, Segond et al. 2021, Bai, Han et al. 2022). Our results are consistent with previous observations documenting the existence of multiple domestication routes in beer and bread yeasts, involving amplification of genes from the *MAL3* cluster (*MAL31*, *MAL32*) and to a lesser extent *IMA1-4* and *MPH2-3* genes or *MAL1* cluster genes (*MAL11*, *MAL12*) (Figure S3F). Yet, a closer investigation of CNVs between genes expected to be in linkage disequilibrium revealed important deviations from the established signatures of domestication.

Because of linkage disequilibrium, relative copy number estimates for *MAL31* and *MAL32* are expected to be strongly correlated. Indeed, we observed strong correlations between CNVs of *MAL31* and hitchhiked genes (*YBR300C*, R^2^ = 0.8098, *p* < 0.0001), as well as between genes within the *MAL1* cluster, although it is more frequently disrupted than amplified within genomes in our dataset (*MAL11 vs. MAL13*, R^2^ = 0.9242, *p* < 0.0001; Figure S3H). In contrast, correlation between CNVs for *MAL31* and *MAL32* genes was incomplete (R^2^ = 0.5957, *p* < 0.0001), while we found no correlation between *MAL11* and *MAL12* CNVs. Notably, relative copy number estimates for *MAL32* and *MAL12* deviated from integer values much more than for *MAL31* (Figure S3H). The common denominators in these discrepancies, *MAL32* and *MAL12*, share > 99.5% sequence identity, implying that misalignment of reads between almost identical *MALx2* paralogues may have confounded the associations between CNVs of linked *MAL* genes. Indeed, clustering reads from *MAL32* and *MAL12* into a *MALx2* (*MAL12, MAL32*) consensus sequence shifted CNVs much closer towards integer values and drastically improved the correlation between *MAL31* and *MALx2* CNVs (R^2^ = 0.8336, *p* < 0.0001; Figures 3A and S3H).

**Figure 3.**
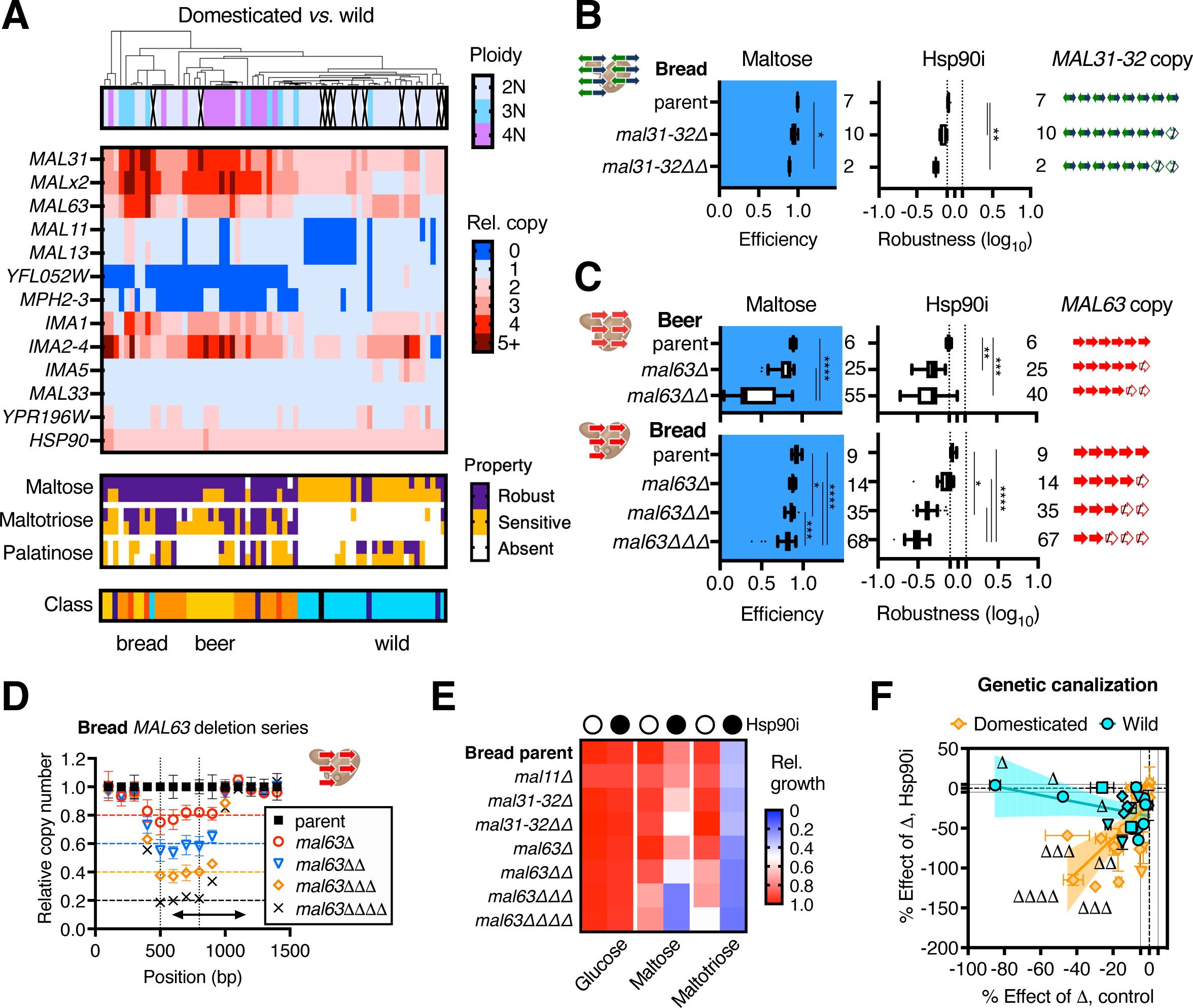
Copy-number signatures of beer and bread domestication canalize industrial traits against Hsp90 stress. (A) Heat map indicates from top to bottom ploidy, CNVs of the indicated genes, robustness of indicated metabolic traits to low- and moderate-level Hsp90 stress, and strain origin. Orange indicates bread strains, yellow indicates beer, red whiskey, purple other fermentation associated, and light blue wild yeasts derived from the gut of wasps. CNVs were determined using CONVICT with clustering set to 90% identity cutoff. (B and C) Genotype schematic and box plots of basal efficiency and robustness of maltose metabolism to moderate-level Hsp90 stress in lineages derived by disruption of (B) *MAL31-MAL32*, and (C) *MAL63* duplications in commercial bread and beer strains. The number of independent biological replicates is indicated next to each plot. (D) CNV analysis of *MAL63* relative copy in bread strain and *mal63* derivatives. 5 *MAL63* gene copies are present in the original parent. Disruption of each *MAL63* copy is expected to reduce the coverage of the deleted region in the gene by 0.2 (1: 5, expected coverage *Δ*: 0.8, *ΔΔ*: 0.6, *ΔΔΔ*: 0.4, and *ΔΔΔΔ*: 0.2) as compared to the corresponding parental strains. Relative coverage is shown at each non-overlapping 100 bp-windows spanning the length of the ORF for 2 – 5 replicate strains per genotype. (E) Heat map of metabolic efficiency for the indicated traits for *MAL* gene disrupted derivatives of commercial bread strain, evaluated under basal and moderate-level Hsp90 stress conditions. (F) Effect of *MAL* gene disruption (Bland-Altman difference) on the efficiency of maltose, maltotriose and turanose metabolism under basal (x-axis) and moderate-level Hsp90 stress conditions (y-axis), and simple linear regression comparisons between wasp gut-derived wild strains (4 different backgrounds: 80, 93, 442, 449) harboring “low” copy numbers of *MAL* genes as compared to domesticated strains (4 different backgrounds: bread, beer, whiskey, rum) with “high” copy numbers (simple linear regression, F = 22.21, DFn = 1, DFd = 44, *p* < 0.0001). Each data point represents a different strain/condition and symbol shape/color indicates different genomic backgrounds. Averages and standard deviations were derived from 2 – 3 independent replicate experiments. Significance between distributions was determined using the Kruskal-Wallis test with Dunn’s multiple comparisons.

A second refinement we implemented in our CNV analysis was inclusion of a template sequence for *MAL63*, a member of the *MALR* family of sugar-inducible transcription factors required for the activation of *MALT* and *MALS* genes (Needleman, Kaback et al. 1984, Brown, Murray et al. 2010). Because *MAL63* is missing from the reference genome, this gene is typically omitted from studies of yeast domestication (Gallone, Steensels et al. 2016, Goncalves, Pontes et al. 2016, Steenwyk and Rokas 2017, Duan, Han et al. 2018, Bigey, Segond et al. 2021). Importantly, the Mal63 transcription factor is an Hsp90 client protein and relies on the Hsp90 chaperone to maintain its function (Bali, Zhang et al. 2003, Ran, Bali et al. 2008, Ran, Gadura et al. 2010). Utilizing a *S. pastorianus MAL63* consensus sequence as a reference, CNV analysis revealed that *MAL63* is amplified across diverse beer and bread strains, in association with the robustness of maltose metabolism against Hsp90 stress and in conjunction with *MAL31* and *MAL32* (*MAL63 vs. MAL31,* R^2^ = 0.7489, *p* < 0.0001, *MAL63 vs. MALx2,* R^2^ = 0.5189, *p* < 0.0001; Figures 3A and S3F). Surprisingly, there was negligible CNV in the *MALR* genes *MAL33* and *YPR196W* (Figure 3A), which have previously been associated with domestication (Goncalves, Pontes et al. 2016, Steenwyk and Rokas 2017, Duan, Han et al. 2018). This discrepancy is explained by accentuated read misalignment between similar *MAL63* and *MALx3* paralogues in the absence of a template reference for *MAL63*. We conclude that amplification of *MAL63*, not *MAL33* or *YPR196W*, is associated with domestication and the robustness of maltose metabolism to Hsp90 stress (Figure 3A). These analyses provide a refined genomic signature of beer and bread domestication. We propose this genomic signature usually involves 3 – 5 additional relative copies of sub-telomeric *MAL* clusters (i.e., *MAL3*, *MAL4*, *MAL6*, *MAL1*) containing maltose metabolism genes and that these genes include *MAL63*, *MAL31*, *MAL32* and, much less frequently, *MAL11*, and *MAL12*.

Importantly, the relative copy number of genes from the *MALx2* family explained a striking 50 – 60% of the genetic variance in the robustness of maltose metabolism to Hsp90 stress (two-tailed Pearson correlation, robustness of maltose metabolism to low-level Hsp90 inhibition (Hsp90i): n = 63, R^2^ = 0.5243, *p* < 0.0001, moderate-level Hsp90i: n = 65, R^2^ = 0.5387, *p* < 0.0001; Figure S3I), and basal maltotriose utilization efficiency (two-tailed Pearson correlation, maltotriose: n = 64, R^2^ = 0.5821, *p* < 0.0001; Figure S3J). The strong association between *MALx2* CNV and robustness of maltose metabolism to Hsp90 stress was not an artifact of ploidy differences between domesticated and wild yeasts or bias towards domesticated strains, because it was reproduced when we limited these comparisons to diploid strains or wild isolates from the gut of wasps (two-tailed Pearson correlation, diploids: low-level Hsp90i: R^2^ = 0.5929, n = 28, *p* < 0.0001, moderate-level Hsp90i: R^2^ = 0.5821, n = 29, *p* < 0.0001, wasp-derived: n = 26, low-level Hsp90i: R^2^ = 0.3486, *p* = 0.0015 and moderate-level Hsp90i: R^2^ = 0.2759, *p* = 0.0059; Figures 3A and S3K-L). Notably, these comparisons revealed a strong threshold effect for *MAL* gene copy number on the robustness of maltose metabolism to Hsp90 stress and basal maltotriose utilization efficiency (Figures S3I-J).

As expected, *MALx2* copy number was weakly correlated with basal maltose utilization efficiency and no significant association was observed with the efficiency of palatinose metabolism (two-tailed Pearson correlation, maltose: n = 64, R^2^ = 0.2855, *p* < 0.0001, palatinose: n = 62, ns; Figure S3M, and S3N). Notably, the copy number of *IMA5*−involved in palatinose metabolism−was positively correlated with the basal efficiency of palatinose metabolism (Pearson correlation, R^2^ = 0.2634, *p* < 0.0001) and weakly associated with the robustness of this trait against Hsp90 stress (Pearson correlation, moderate-level Hsp90i: R^2^ = 0.1337, *p* = 0.0334). These observations imply that *MAL* gene duplications influence the robustness of metabolic traits to Hsp90 stress. Additionally, we observed that certain beer strains (4 lager yeasts and 1 ale strain) harbored 1 extra relative copy of *HSP90*. However, this did not explain the increased robustness of metabolic traits to Hsp90 stress, even though *HSP90* overexpression is able to confer such increased robustness (Figures 3A and S1J). On the other hand, we found significant associations between the robustness of maltose metabolism to Hsp90 stress and amplifications or deletions in genes with no or only secondary functions in maltose and maltotriose metabolism *(IMA1*, *IMA2-5*, *MPH2-3*, *MAL13*, *YFL052W* (Figures 3A and S3F). Although CNVs of these genes have been previously associated with yeast domestication (Gallone, Steensels et al. 2016, Goncalves, Pontes et al. 2016, Steenwyk and Rokas 2017, Bigey, Segond et al. 2021), the above observations suggest caution in interpreting the functional significance of CNVs based on associations alone. Nevertheless, the findings presented above demonstrate that the robustness of traits to Hsp90 stress is genetically determined and provide refined genomic signatures of yeast domestication to help in establishing causal relationships between *MAL* gene duplications and canalization against Hsp90 stress.

### Redundant gene duplications canalize industrial traits against Hsp90 stress

To investigate a causal relationship between *MAL* gene duplications and trait canalization, we determined the effect of *MAL* gene disruptions on the sensitivity of maltose metabolism to Hsp90 inhibition in isogenic derivatives of diverse industrial and wild yeasts harboring different numbers of *MAL* gene copies. To maximize the efficiency and specificity of gene editing while accounting for possible differences in transformation efficiency and homologous recombination between strains, we utilized selection cassettes harboring > 500-bp long homology arms spanning regions at the 5’ and 3’ ends of the targeted *MAL* gene ORFs (Figure S3O).

We generated a series of *MAL* mutations in diverse industrial (beer, bread, whiskey, rum) and wild (derived from the gut of wasps) strains. Because *MAL31* and *MAL32* genes are typically co-inherited, we disrupted both genes in one step, also deleting the bidirectional promoter that drives their Mal63-dependent expression. Notably, disruption of a single copy of *MAL31* and *MAL32* had no obvious effect on basal maltose utilization efficiency (left panel; Figure 3B). Maltose metabolism was similarly robust to *MAL63* disruptions across diverse industrial strains (left panels; Figures 3C, S3P and S3R). We observed some small effects on basal metabolism only after 2 or 3 *MAL* gene disruption steps (Figures 3B, 3C and S3P-R), while several industrial strains tolerated up to 3 or 4 *MAL* gene disruptions with negligible effects on basal maltose utilization efficiency (Figures 3C, S3Q and S3R).

We confirmed the deletions in 38 strains derived from 6 parental strains by whole genome sequencing and CNV analysis. As expected, each derivative strain yielded proportionately reduced coverage of the disrupted *MAL* gene within the deleted region as compared to the cognate parental strain (Figures 3D and S3S). Additionally, the derivative strains also harbored one copy of the selection cassette (*natMX, hphMX, kanMX*) for each disrupted *MAL* gene. Notably, both maltose and maltotriose metabolism were highly robust against *MAL* gene disruptions in domesticated yeasts (Figure 3E). The striking robustness of these traits was unexpected given that *MAL* gene duplications are thought to support basal efficiency of maltose and maltotriose metabolism (Duval, Alves et al. 2010, Gallone, Steensels et al. 2016, Bigey, Segond et al. 2021, De Chiara, Barre et al. 2022). Further arguing against this conventional view, metabolic traits were substantially more robust to mutations in domesticated yeasts as compared to wild isolates. Disruption of a single *MAL* gene reduced the basal efficiency of maltose, maltotriose and turanose metabolism across diverse wild strains, whereas disruption of two or more genes was required to affect these traits across domesticated strains, which reflects the lower number of *MAL* genes in the genome of non-domesticated strains (Figure 3F). Overall, the results demonstrate that industrially relevant metabolic traits experienced genetic canalization during yeast domestication by virtue of redundant *MAL* gene duplications.

Next, we investigated the role *MAL* gene duplications play in the robustness and canalization of industrially relevant metabolic traits to Hsp90 stress. Strikingly, disruption of either *MAL31*-*MAL32* or *MAL63* genes profoundly reduced the robustness of maltose and maltotriose metabolism to Hsp90 inhibition across diverse industrial strains (Figures 3B-F and S3P-R). Disruption of *MAL* gene copies reduced trait robustness reproducibly across independent biological replicates of derivative strains (n = 10–67), suggesting that the disrupted genes encoded for functional proteins (Figures 3B-C, and S3P-R). These findings demonstrate that redundant *MAL* gene duplications which give rise to genomic signatures of yeast domestication canalize industrially relevant metabolic traits (maltose and maltotriose metabolism) in industrial strains against Hsp90 stress. Given the strong adaptive role of *MAL* gene duplications during yeast domestication, our results suggest that the profound canalization of industrial traits against Hsp90 stress evolved as an adaptation to maltose-rich domestication environments (adaptive canalization).

### Hsp90 stress from proteotoxic metabolite de-canalizes industrial traits

Validation of a model for yeast domestication by adaptive canalization of maltose metabolism against Hsp90 stress relies on the identification of biotic or abiotic environmental stressors that could exert persistent Hsp90 stress on industrial yeasts and would be associated with industrial fermentations. Beer, bread, and whiskey strains must endure diverse fermentation-associated stressors, several of which are known to be proteotoxic in yeast and other systems (Herskovits, Gadegbeku et al. 1970, Cowen and Lindquist 2005, Rohner, Jarosz et al. 2013, Karras, Yi et al. 2017, Alford and Brandman 2018, Voordeckers, Colding et al. 2020).

To test the validity of an adaptive canalization model for yeast domestication, we first tested stressors associated with industrial niches (acidosis, elevated temperatures, salinity, alcohols) by comparing their effects on maltose metabolism to the effects of Hsp90 inhibition across diverse wild-derived *S. cerevisiae* strains of the same ploidy. Most exposures perturbed maltose utilization in a strain-specific way, without proportional effects on growth in glucose. Among the conditions tested, heat stress (37°C) and ethanol exposure closely reproduced the effects of Hsp90 inhibition on maltose utilization (Figure S4A). In addition to inhibiting maltose metabolism, ethanol exposure also inhibited maltotriose and palatinose metabolism in a strain-specific way across diverse domesticated and wild yeasts, without imposing proportional growth constraints (Figure 4A). Notably, effects were observed at ethanol concentrations well within the range at which the metabolite is found in most industrial fermentation niches (4 – 12%, R^2^ = 0.813, *p* < 0.0001; Figure S4B). Moderate ethanol stress (6%, v/v) closely reproduced the effects of low-level Hsp90 inhibition on the efficiency of metabolic traits (Figure 4B). In agreement with ethanol serving as a major selection agent in the environmental canalization model, maltose metabolism was more robust against ethanol stress across domesticated yeasts in maltose-rich niches as compared to wild strains or domesticated strains from maltose-poor niches (Figures S4C and S4D). Moreover, beer, bread and 3 whiskey strains utilized maltose substantially more robustly under moderate ethanol stress as compared to all other groups, and also under high Hsp90 stress elicited by the combined effect of ethanol exposure and moderate-level Hsp90 inhibition (ethanol 8% v/v + radicicol 10 μM; Figures 4C and S4F). Additionally, the robustness of maltotriose metabolism to ethanol exposure and low or high Hsp90 stress was associated with domestication niche (Figures S4C and S4F). We found no significant associations with robustness to ethanol stress for palatinose metabolism, except for the wild isolates mirroring our observations with Hsp90 inhibition as mentioned above (compare Figure S4C to S2J, Figure S4D to S2F, and Figure S4E to Figure 2E). The effects of ethanol exposure on metabolic efficiency were highly reproducible across technical and biological replicate experiments, similar to basal and Hsp90 inhibition conditions (Figure S4G). These findings are consistent with the adaptive canalization hypothesis of yeast domestication and demonstrate that ethanol exposure impairs Hsp90-dependent metabolic traits at concentrations consistent with those found in maltose-rich industrial niches.

**Figure 4.**
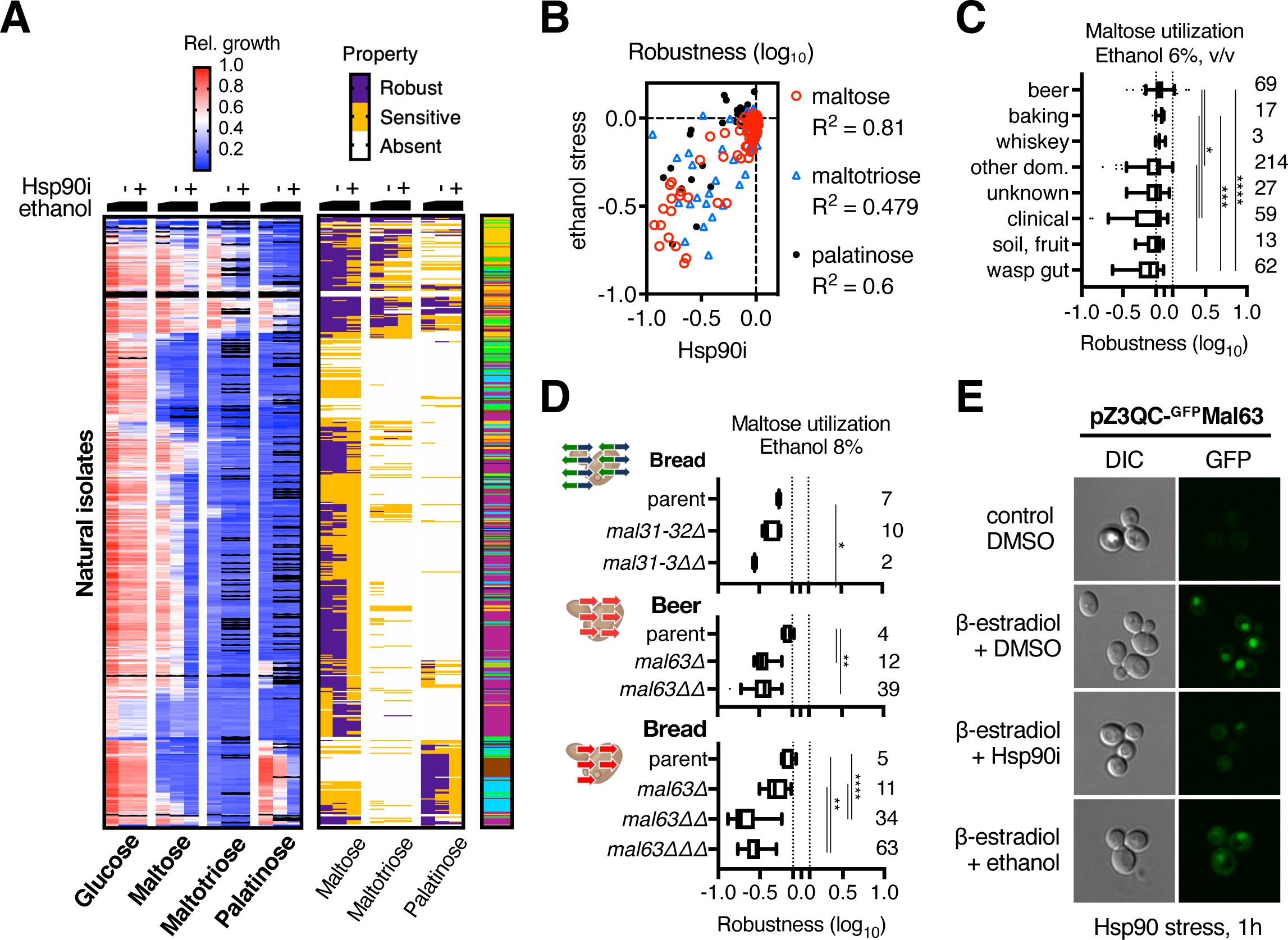
Ethanol exposure de-canalizes traits by eliciting Hsp90 stress. (A) Heat map of relative growth and trait robustness estimates for 534 natural *Saccharomyces* isolates evaluated under different levels of ethanol stress, from left to right: 6%, 8% with DMSO and 8% with radicicol (10 μM). Strain origin is indicated in the color key in Figure 2A. (B) Correlation between the robustness to low-level Hsp90 inhibition (radicicol, 4 μM) and ethanol stress (6%, v/v) for maltose (Pearson R^2^ = 0.8095, *p* < 0.0001, n = 88), maltotriose (Pearson R^2^ = 0.4789, *p* < 0.0001, n = 51), and palatinose metabolism (Pearson R^2^ = 0.5986, *p* < 0.0001, n = 35) evaluated across diverse domesticated and wild yeasts (described in Figure S3C). (C) Box plots of robustness to ethanol stress (6%, v/v) evaluated for maltose metabolism across all natural isolates indicated in Figure 4A and parsed by strain origin. (D) Genotype schematic and box plots of robustness of maltose metabolism against ethanol stress (8%, v/v) across *mal31-32* and *mal63* disrupted derivatives of industrial bread and beer strains. (E) Confocal microscopy of BY4741 cells expressing GFP-Mal63 in the presence of β-estradiol (100 nM) under basal and Hsp90 stress conditions (1 hour exposure with radicicol, 10 μM, *vs.* ethanol, 6%, v/v). Values and standard deviations were derived from 2 – 3 independent biological replicates and 2 independent experiments. Significance between distributions was determined using the Kruskal-Wallis test with Dunn’s multiple comparisons.

Next, we evaluated the relationship between *MAL* gene copy number and the robustness of metabolic traits against ethanol stress. As expected, the robustness of maltose metabolism to ethanol exposure was strongly associated with an increased copy number of *MAL* genes, whereas no significant associations were observed for palatinose utilization genes, *HSP90 (HSC82, HSP82)*, housekeeping genes (*UBC6*) or control genes (two-tailed Pearson correlation, *MALx2* R^2^ = 0.4739, *p* < 0.0001; Figures S4H and S4I). Moreover, sequential disruption of *MAL* genes increased the sensitivity of maltose utilization to ethanol exposure across diverse industrial strains (Figure 4D), suggesting that ethanol exposure reveals Hsp90-buffered CNVs of *MAL* genes. Additionally, we observed a dose-dependent threshold relationship between the level of Hsp90 stress and the number of *MALx2* copies required for robust maltose utilization (robustness cut-off −0.25, ethanol 6%: 2.1 *MALx2* copies, ethanol 8%: 2.65 *MALx2* copies, ethanol 8% + Hsp90i: 4.2 *MALx2* copies, n = 65, *p* < 0.0001; Figure S4J). We observed similar relationships between ethanol concentration and the copy number of *MAL31* (6%: 2.11, 8%: 2.87, ethanol + Hsp90i: 4.18, n = 65) and *MAL63* (6%: 1.0, 8%: 1.8, ethanol + Hsp90i: 3.88, n = 65). Furthermore, ethanol exposure led to profound trait de-canalization, exposing inherent Hsp90 dependencies in robust metabolic traits across diverse domesticated and wild yeasts (Figure 4A). These data demonstrate that the Hsp90 dependencies of maltose and maltotriose metabolism were conserved during yeast domestication. Mal63 is most likely the only Hsp90-dependent component in the maltose and maltotriose metabolism pathways .

In support of Mal63 being a major point of fragility in these maltose-utilization pathways, moderate ethanol stress drove GFP-tagged Mal63 out of the nucleus and into the cytoplasm and also depleted the protein in a time-dependent fashion, phenocopying Hsp90 inhibition (Figures 4E and S4K). Ethanol did not affect the stability of GFP or Hsp90 expression over the duration of these experiments (Figure S4K). As a control, we confirmed that ^GFP^Mal63 supported maltose and maltotriose metabolism comparably to the untagged Mal63 protein (Figure S4L). Notably, overexpression of Mal63 protein rendered both traits robust to Hsp90 stress, without affecting the basal efficiency or robustness of palatinose utilization (Figure S4L). On the other hand, overexpression of an unrelated MalR protein, Mal13—required for palatinose utilization— increased the robustness of palatinose metabolism to Hsp90 impairment, without affecting maltose or maltotriose metabolism under basal or Hsp90 inhibition conditions (Figure S4L). Mal63 is most likely the only MalR transcription factor driving maltose and maltotriose metabolism. This is in line with our findings that disruption of sufficient *MAL63* copies selectively blocked these and other pathways (turanose metabolism) across diverse genomic backgrounds (Figures 3C, 3E, 3F, S3P and S4M). Taken together, these findings demonstrate that ethanol exposure compromises Hsp90 chaperone capacity, thereby uncovering cryptic, Hsp90-buffered CNVs in metabolism (de-canalization), and therefore satisfies a major criterion for an important role in adaptive canalization.

### Ethanol exposure transforms the adaptive value of metabolic gene duplications

Identification of ecologically relevant Hsp90 stressors interacting with *MAL* gene duplications does not necessarily prove that environmental canalization is adaptive, especially because adaptive canalization is believed to be limited by alternative scenarios involving selection against biological noise (Gavrilets and Hastings 1994, Burga, Casanueva et al. 2011), selection against lethals (Nowak, Boerlijst et al. 1997, Siegal and Bergman 2002, Stearns 2002), pleiotropy (by-product of other adaptations) (Nowak, Boerlijst et al. 1997, Wagner, Booth et al. 1997, Milloz, Duveau et al. 2008, Bauer, Li et al. 2015) and genetic drift (Payne and Wagner 2019). Distinguishing between these canalization scenarios is challenging, especially given that a role for Hsp90 has been proposed in each of these models. For example, stochastic noise in Hsp90 expression between single cells has been shown to affect the penetrance of genetic variations in *C. elegans* (Burga, Casanueva et al. 2011). In our system, a selection-against-noise scenario would predict that *MAL* gene duplications confer a fitness advantage to industrial yeasts growing in maltose even under basal conditions by canalizing maltose metabolism against biological noise, such as stochastic fluctuations in gene expression (i.e., Hsp90 level, Mal63 stability). Likewise, a selection-against-lethals model would predict that *MAL* gene duplications increase fitness by rendering maltose metabolism robust against detrimental mutations (stochastic or induced). In contrast, a pleiotropy-based model would require that canalization result from selection for a pleiotropic character that indirectly reinforces trait robustness against perturbations. According to such a model, *MAL* gene duplications should confer a strong effect on fitness in the presence of selection for the hypothetical pleiotropic trait that also favors canalization of maltose utilization. Finally, the adaptive canalization model predicts that *MAL* gene duplications would confer a strong fitness advantage to industrial yeasts in maltose-rich environments, but only in the presence of Hsp90 stress.

To help distinguish which mechanism or combination thereof may have shaped the profound genetic and environmental canalization of maltose metabolism that we observed across industrial yeasts, we adopted a competition-based approach (Wilkins 1997). We determined the impact of *MAL* gene duplications on the relative fitness of isogenic, disrupted derivatives of an industrial bread yeast strain (Figure 5A) competing in maltose-rich environments. Because use of bulk growth parameters as proxies of relative fitness could lead false inferences about the strength of selection (Chevin 2011), we labelled strains by stably expressing different fluorescent proteins under the control of a strong constitutive (GPD) promoter from the *HIS3* genomic locus. Labeling cells with msfGFP and mRuby3 produced populations in which 99.7% of the cells were fluorescent, their median fluorescence being 30– 100-fold above background (Figures S5A and S5B). Lineages of labelled strains retained fluorescence under diverse conditions, including ethanol stress, presenting the same morphology and growth parameters as the parental strain (Figures 5B, 5C and S5A-C). Next, we mixed the labelled strains in 1:1, green *versus* red, cell ratio and allowed them to compete in the presence of maltose. We observed a small selective advantage for the parent strain as compared to the single *MAL31-32* copy disrupted derivative, and this effect was markedly increased upon Hsp90 inhibition independently of fluorescent marker configuration (Figure S5D). Yet, the parental strain reached fixation within a single competition round, which precluded quantitative assessment of relative fitness effects. Hence, we repeated this experiment using a 1:1,000 dilution ratio where the parent represents 0.1% of the initial cell population, propagating cultures through sequential bottlenecks. Hsp90 inhibition drastically exacerbated (by ∼20-fold) the relative fitness of *MAL31-32* gene duplications, which was weak under basal conditions (Figure S5E). Swapping the fluorescent marker between strains did not affect the relative fitness of the parental strain (Figure S5F). As expected, parental strains did not out-compete disrupted derivatives in glucose media (Figures S5E and S5F). However, the effect of Hsp90 inhibition was again too strong to quantitatively evaluate the strength of selection it exerted.

**Figure 5.**
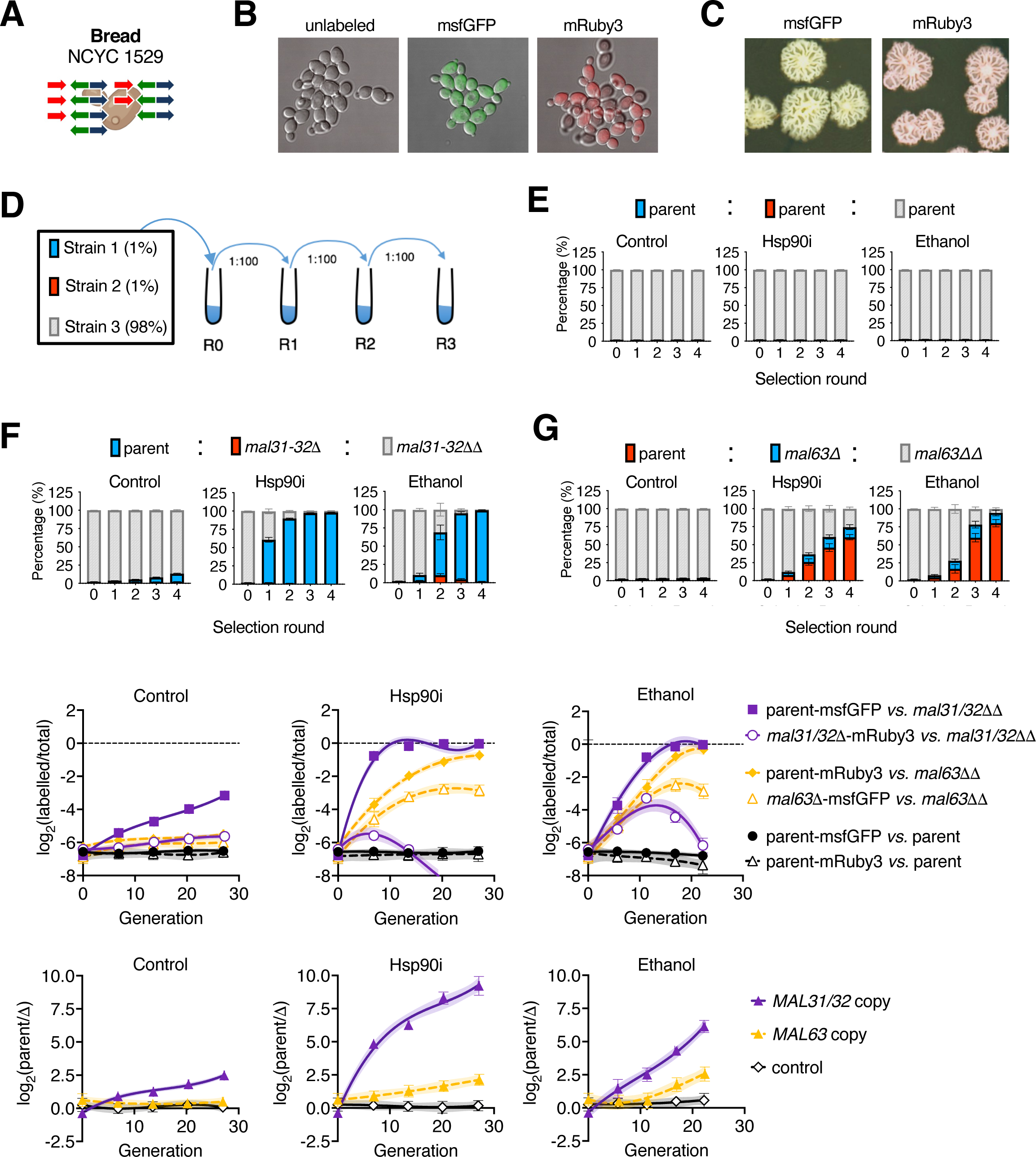
Ethanol selects for redundant gene duplications conducive to adaptive canalization. (A) Schematic of genotype of parental commercial bread strain, which harbors 7 *MAL31-32* and 5 *MAL63* copies. (B) Confocal microscopy of fluorescently labelled derivatives forming micro-colonies in SM media. (C) Image of fluorescent colonies observed under visible light after growth on YPD agar plates for 9 days at room temperature. (D) Schematic of selection setup for competition between isogenic derivative strains. Dilutions and timing between each selection round, R, vary across experiments as indicated (main text and STAR methods). (E-G) Cell cytometry analysis of 1:1:98 mixed populations passing over sequential broad bottlenecks (1:100) and allowed to expand in YPM under basal, moderate-level Hsp90 inhibition (radicicol, 10 μM), or ethanol stress (8%, v/v) conditions, using the following strain combinations: (E) parental strains competing against each other, (F) single (mRuby3) and double *mal31-32* disruption lineages competing against the parent (msfGFP), and (G) single (msfGFP) and double *mal63* disruption lineages competing against the parent (mRuby3). (H and I) Log2 ratios of genotypic frequencies plotted over the number of generations under basal conditions (left panels) evaluated against (H) non-labelled double disrupted derivatives (*mal31-32ΔΔ*, *mal63ΔΔ*), and (I) labelled single disrupted (*mal31-32Δ*-msfGFP, *mal63Δ*-mRuby3) strains, competing against labelled parent derivatives, as indicated above. Values and standard deviations were derived from 4 independent technical replicates. Significance between distributions was determined using the Kruskal-Wallis test with Dunn’s multiple comparisons.

We overcame this limitation by increasing the effective population size during inoculation to reduce the number of generations per competition round from ∼10.5 to ∼6.6, which more closely reflects standard back-slopping practices in traditional fermentations (Whittington, Dagher et al. 2019, De Vuyst, Comasio et al. 2021). Moreover, to simulate genetic variation in ancestral pre-domestication communities, we competed labeled parental and single *MAL* gene-disrupted derivative strains with unlabeled double-disrupted derivatives at a 1:1:98 ratio (Figure 5D). *MAL* gene duplications conferred strong selective advantage only under conditions of Hsp90 stress in both *mal31-32* and *mal63* mutant lineages (Figures 5E-G). Hsp90 inhibition increased the effects of *MAL* duplications on relative fitness by > 20—30-fold (Figure 5H). In fact, a single *MAL31-32* or *MAL63* copy conferred fitness advantage > 20-fold in maltose under Hsp90i conditions, when comparing parental lineages to single *MAL* copy-disrupted derivatives (Figures 5H and 5I). Under Hsp90 stress, strains with extra *MAL* gene copies readily out-competed doubly disrupted derivatives, and single disrupted strains were eventually out-competed by the parental strains (Figures 5H and 5I). Notably, the effect of disruption of two *MAL63* copies on relative fitness in maltose under Hsp90i conditions was comparable to disruption of a single *MAL31-32* copy (Figures 5H and 5I).

Although we observed effects of single and double *MAL31-32* disruption on relative fitness in maltose under basal conditions, these effects paled in comparison to the fitness advantage *MAL31-32* duplications conferred under Hsp90i conditions (Figures 5H and 5I). Single disruption of *MAL63* had no effect on relative fitness in maltose under basal conditions, while we observed a small effect for two *MAL63* disruptions (basal: *s*_1x*MAL63*_ = −0.4243% ± 0.7555%, *s*_2x*MAL63*_ = +2.525% ± 0.3202%; Figure 5, G-I). In comparing observed effects in this setting, we estimated that it would take > 150 generations for two *MAL63* copies to reach 50% frequency in maltose under basal conditions, as opposed to merely 17 generations under conditions of Hsp90 inhibition (moderate Hsp90i: *s*_2x*MAL63*_ = +22.2% ± 0.049%). Importantly, ethanol stress replicated the effects of Hsp90 inhibition in this assay (Figures 5E-I). Results were reproducible using independent isogenic strains and using independent methods for measuring relative fitness (cell cytometry *vs.* colony counting on plates; Figures S5G-I). In all experiments, ethanol stress revealed strong cryptic fitness effects for redundant *MAL* gene duplications in maltose-rich media. We also reproduced the interaction between *MAL* gene copy number and ethanol exposure on relative fitness in wort and sourdough media, even at low concentrations of ethanol (5%, v/v) (Figures S5J-L). Notably, the selection coefficient of extra *MAL* gene copies was on the order of 35 – 72% in maltose-rich media under ethanol stress or Hsp90i conditions. Taken together, results from competition experiments argue against a major role for biological noise or basal metabolic flux in driving the canalization of maltose metabolism. Further, they demonstrate that ethanol exerts strong selective pressure for the *MAL* gene duplications characteristic of domesticated yeasts, thus favoring the adaptive model of yeast domestication by environmental canalization of maltose metabolism.

Next, we examined potential canalization of maltose metabolism by selection-against-lethals. In this model, extra *MAL* gene copies buffer against deleterious mutations acquired spontaneously or in response to stress. Industrial yeasts can clearly undergo rearrangements in response to stress, including at preexisting sites of CNVs (Hull, Cruz et al. 2017). Hsp90i may accelerate this process by increasing the rate at which new mutations emerge (James, Usher et al. 2008, Dunn, Paulish et al. 2013, Zabinsky, Mason et al. 2019). Notably, ethanol exposure can also increase mutation rates by causing DNA replication stress (Voordeckers, Colding et al. 2020). However, ethanol’s mutagenic effect is modest (2.6 – 3.5-fold under 6%, v/v) and would not be expected to drive strong selection against lethals within evolutionary periods relevant to industrial fermentations−up to 150 generations per year for modern beer strains (Gallone, Steensels et al. 2016). On the other hand, ethanol exposure rapidly selected for canalizing *MAL* gene copies within < 17 generations (Figures 5E-I and S5D-K), which provides insufficient time for new mutations to shape trait robustness through negative (or positive) selection. These observations disfavor a model of canalization involving selection for or against new mutations and reveal that ethanol exerts strong selection pressure for trait robustness by revealing preexisting Hsp90-buffered genetic variation within the population.

Our data also provide strong evidence against a model of canalization based on pleiotropy (by-product hypothesis). *MAL11* encodes a promiscuous permease with broad specificity across diverse sugars (Alves, Herberts et al. 2008, Brown, Murray et al. 2010). However, *MAL11* is more frequently lost or inactivated than it is amplified across domesticated strains (Vidgren, Ruohonen et al. 2005, Gallone, Steensels et al. 2016, Gallone, Mertens et al. 2018), and its association with robustness of maltose utilization to Hsp90 stress was driven by reduced *MAL11* copy number (two-tailed Pearson correlation, low-level Hsp90i R^2^ = 0.1943, *p* = 0.0003, n = 63, moderate-level Hsp90i R^2^ = 0.2782, *p* < 0.0001, n = 63, *vs.* ns after removal of 8 genomes completely lacking *MAL11*; Figure 3A). Another pleiotropic gene family in the pathway, *MALx2*, encodes promiscuous enzymes that hydrolyze maltotriose, turanose, and sucrose in addition to maltose (Brown, Murray et al. 2010). If canalization of maltose metabolism was a pleiotropic effect of selection for efficient basal maltotriose, turanose, or sucrose metabolism, then disruption of *MAL31-32* genes would be expected to drastically reduce the basal efficiency of various metabolic traits. In contrast, disruption of two *MAL31-32* copies in an industrial strain barely affected the basal efficiency of maltose, maltotriose, turanose, and sucrose metabolism (Figure 3E and Figure S4M). Moreover, *MAL* gene disruptions had negligible basal fitness effects in wort and sourdough media, which contain large amounts of maltotriose (Figures S5J and S5K). The same genetic perturbations drastically reduced the robustness of these traits under Hsp90 inhibition and ethanol exposure, except for sucrose metabolism which remained robust against all these perturbations (Figures 3B, 3C, 3E, 3F, 4D, S3P-R and S4M). Furthermore, under Hsp90 stress conditions, *MAL* gene duplications conferred similar competitive benefits to industrial yeasts in wort and liquid sourdough as compared to media containing maltose as sole carbon source (Figures 5F-I and S5D-L). Notably, maltotriose in wort and dough begins to be utilized during industrial fermentations after most of the maltose has been converted to ethanol and the yeast has stopped growing (Patel and Ingledew 1973, Zheng, D’Amore et al. 1994, Londesborough 2001), suggesting that maltotriose metabolism must be robust to ethanol stress to confer a fitness advantage in industrial niches. Indeed, maltotriose utilization was highly robust to moderate ethanol stress (6%, v/v) across beer, bread and whiskey yeasts as compared to strains from other ecological niches (Two-tailed Mann-Whitney test, *p* < 0.0001, n_bbw_ = 74 *vs.* n_other_ *=* 107).

Taken together, our findings argue against various non-adaptive models of canalization of industrial maltose metabolism, namely evolution by genetic drift, selection on basal metabolic flux, selection against biological noise, selection against lethals, and pleiotropy. Instead, they provide strong support for a model of environmental (and congruent genetic) canalization in which the profound robustness of industrial traits to Hsp90 impairment is an adaptation to niche-related Hsp90 stress likely exerted by ethanol in maltose-rich domestication niches (Figure 6).

**Figure 6.**
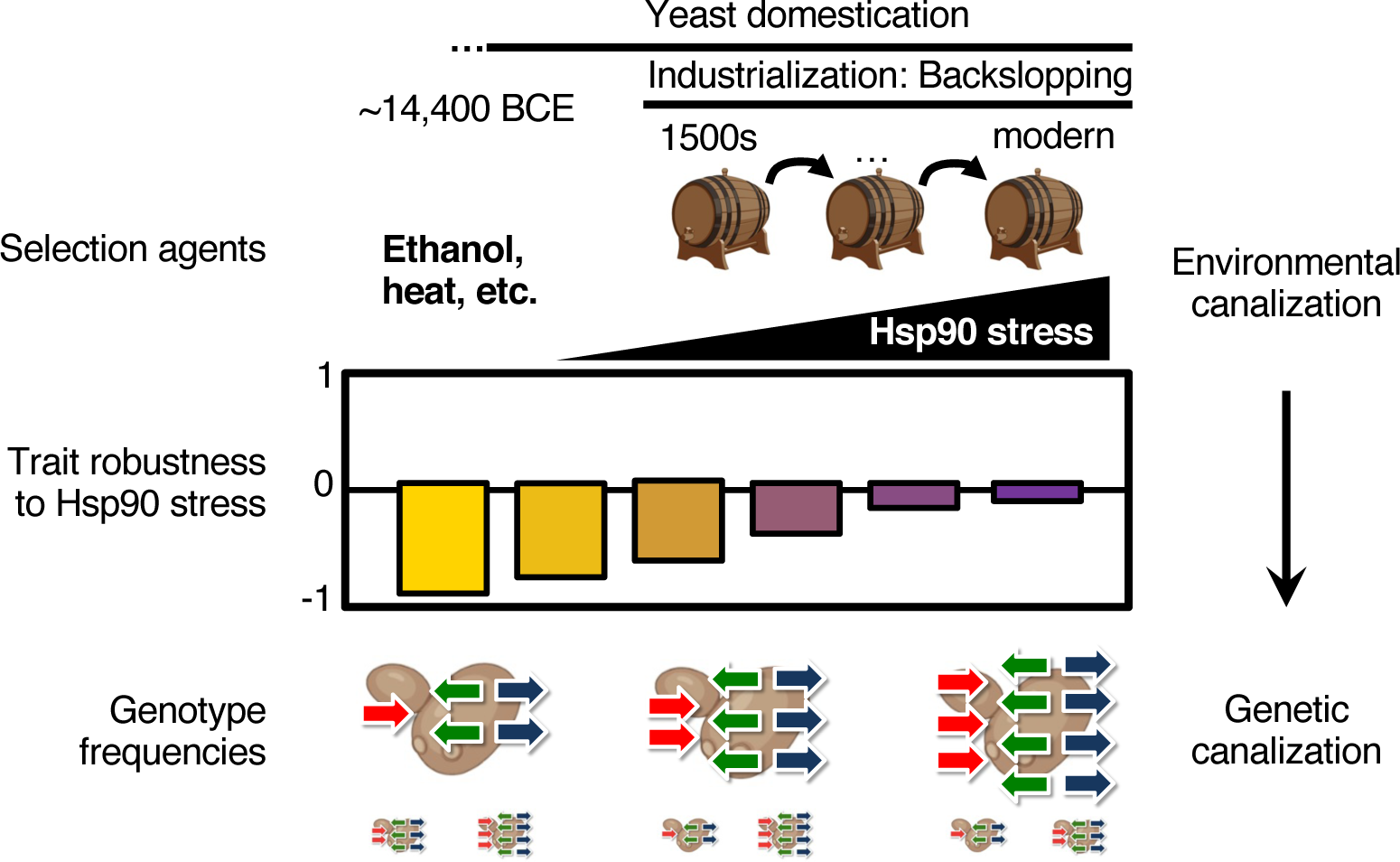
Canalization of industrial traits against niche-related Hsp90 stress enables yeast domestication. Model of environmental canalization and congruent genetic canalization of Hsp90-dependent industrial traits. Hsp90-potentiated metabolic traits in yeast are inherently sensitive to proteotoxic stressors in the environment of the cell. Ethanol accumulation during fermentation derails metabolic traits by causing Hsp90 stress. Neutral evolution over millions of years enabled the accumulation of cryptic genetic variation in ancestral yeast populations which allowed for brewing and bread making before the 1500s. Industrialization and freezing of live beer and bread strains enabled selection for pre-existing *MAL* gene duplications that impart robustness to economically important metabolic traits against ethanol stress. Size of filled arrows indicates frequency of genotypes harboring the indicated *MAL* gene duplications in the ancestral and modern industrial populations of beer and bread yeasts.

## Discussion

Since first proposed over 70 years ago, the concept of canalization has received widespread consideration, but its potential practical applications remain underdeveloped (Dewitt, Sih et al. 1998, West-Eberhard 2005, Koonin and Wolf 2009, Fusco and Minelli 2010, Hull, Cruz et al. 2017, Kelly 2019). Major reasons may be that the mechanisms underlying this fundamental biological phenomenon are incompletely understood and have been traditionally studied in artificial settings that do not recapitulate both the natural ecology and fitness effects of these mechanisms or do not allow testing alternative hypotheses to the one tested (Wagner, Booth et al. 1997, Rutherford and Lindquist 1998, Cowen and Lindquist 2005, Wagner 2005, Rohner, Jarosz et al. 2013, Keane, Toft et al. 2014, Felix and Barkoulas 2015, Geiler-Samerotte, Zhu et al. 2016, Naseeb, Ames et al. 2017, Hallgrimsson, Green et al. 2019, Payne and Wagner 2019, Crespi, Burnap et al. 2022). Here, we make important progress towards addressing these shortcomings by identifying both the ecological stressors and robustness mechanisms driving adaptive canalization in a model eukaryote that has been evolving within anthropic niches of great economic importance.

Budding yeasts are connected with much of human history but the mechanisms underlying the species unrivaled biological success and profound socioeconomic impact remain incompletely understood (Steensels, Gallone et al. 2019, Lahue, Madden et al. 2020, Bai, Han et al. 2022). Over the last few centuries, *S. cerevisiae* underwent strong selection for the ability to produce ethanol in sugar-rich anthropic niches. Alcoholic fermentation of maltose and maltotriose relies on well-understood and genetically similar metabolic pathways critical to produce diverse foods and beverages (i.e., beer, bread, whiskey) and these pathways have markedly evolved during yeast domestication (Steensels, Gallone et al. 2019, Lahue, Madden et al. 2020). However, the adaptive value of these evolved pathways in beer and bread domestication has eluded experimental demonstration (Goddard and Greig 2015, Steenwyk and Rokas 2018). Now, we show that metabolic gene duplications associated with yeast domestication reinforce desirable industrial traits against compromise of Hsp90 chaperone function by stressors prevalent in maltose-rich industrial niches. Ethanol exposure, at relevant concentrations, disrupts key Hsp90-dependent nodes of metabolism and exerts strong selective pressure for gene duplications characteristic of beer and bread yeast domestication. Results from competition assays demonstrate the adaptive value of such canalization and refute alternative “by-product” or “intrinsic” hypotheses. This work identifies maltose utilization in industrial bread and beer yeasts as a long-sought example of *bona fide* adaptive environmental canalization in eukaryotes.

Identification of Hsp90-dependent robustness as a central factor enabling yeast domestication helps answer long-standing questions that have confronted the study of canalization. First, it resolves the controversy around the role of redundancy as a mechanism of canalization in nature (Wilkins 1997, Siegal and Bergman 2002, de Visser, Hermisson et al. 2003, Wagner 2005, Keane, Toft et al. 2014, Hallgrimsson, Green et al. 2019). We show that redundant *MAL* gene duplications are causally responsible for a striking 60% or more of genetic variance in the robustness of maltose metabolism to Hsp90 stress. Canalization of maltose metabolism by redundancy involves CNVs of genes encoding for both the Hsp90-dependent transcription factor, Mal63, and its targets. Hence, maltose metabolism serves as an ecologically relevant paradigm for canalization by redundancy.

Second, this work uncovers a relationship between redundancy and distributed robustness in canalization. Removing layers of redundancy from canalized metabolic traits exposes a cryptic layer of distributed robustness provided by Hsp90’s buffering capacity. This distributed robustness layer canalizes maltose metabolism across diverse industrial beer and bread yeasts against ethanol stress and disruption of *MAL* genes. These findings reconcile two schools of thought on canalization (Siegal and Bergman 2002, de Visser, Hermisson et al. 2003, Wagner 2005) by demonstrating pervasive epistasis between seemingly disparate mechanisms of robustness.

Importantly, we find that different robustness mechanisms differ in their adaptive value. Results from competition assays demonstrate that the adaptive function of redundancy conferred by *MAL* gene duplications resides in the stabilization of maltose and maltotriose metabolism in the face of niche-related Hsp90 stress. Genetic canalization of industrial traits emerged congruently from their adaptive environmental canalization, like for RNA folding (Ancel and Fontana 2000, Szollosi and Derenyi 2009). On the other hand, distributed robustness mechanisms appear intrinsic to metabolic traits; these mechanisms include Hsp90 chaperone capacity and other stress avoidance pathways. The striking preference for redundancy over alternative mechanisms of canalization may reflect evolutionary constraints on the latter, such as the fitness cost of chaperone overexpression (Sabater-Munoz, Prats-Escriche et al. 2015, Cote-Hammarlof, Fragata et al. 2021). Even redundancy is subject to constraints (Naseeb, Ames et al. 2017). For example, the ancient positioning of *MAL* genes into unlinked sub-telomeric clusters may have fostered asymmetric *MAL* gene involvement in canalization. Hence, physical, ecological and evolutionary constraints shape the adaptive value of redundancy and distributed robustness.

It is important to emphasize that buffering mechanisms can have adaptive functions even if they did not evolve adaptively (Wagner and Zhang 2011). This is illustrated here by the correlated effects of redundant *MAL* gene duplications on maltose and maltotriose metabolism; *MAL63* supports both traits by pleiotropy while *MAL32* supports maltose metabolism by linkage. Considering the relative timing of maltose and maltotriose utilization, the ability to utilize maltotriose as sole carbon source may have evolved as pleiotropic byproduct of selection for robustness of maltose metabolism to Hsp90 stress in industrial niches. Another example is the highly contextual roles Hsp90 plays in evolution as genetic potentiator, buffer, and capacitor, although the chaperone evolved to buffer proteotoxic environmental stress (Geiler-Samerotte, Zhu et al. 2016, Iyengar and Wagner 2022, Iyengar and Wagner 2022). In the context of peripheral carbon metabolism, Hsp90 acts as a potentiator for genetic variation in diverse metabolic pathways, suggesting an adaptive role in the diversification of yeast metabolism. As expected for potentiated traits, canalization is a recent event in the evolutionary history of metabolic traits. Indeed, maltose metabolism is canalized against Hsp90 stress across lager beer yeasts but is sensitive in strains related to the closest ancestors of lager beer yeasts (*S. eubayanus*, *S. uvarum*). In contrast, palatinose utilization remained sensitive to Hsp90 stress across yeast species. The Hsp90-potentiation of metabolism for over 100 million years fostered the accumulation of *MAL* gene variation which allowed yeast adaptation in maltose-rich anthropic niches especially during the last 500 years.

In addition to acting as potentiator, Hsp90 also acts as a buffer for variations in *MAL* gene copy number. This observation adds structural variations to the spectrum of genetic alterations Hsp90 interacts with. Yet, this Hsp90 function is not adaptive at least during the timescales we examined. Notably, Hsp90 can act both as a potentiator and a buffer of variation with respect to the same gene (*MAL63*) in the same individual organism; these functions are not antagonistic. In fact, they originate from the same biochemical mechanism that is the Hsp90-dependent maturation and nuclear localization of the maltose-responsive Mal63 transcription factor which shares striking similarities with other ligand activated, obligate, Hsp90 clients (Bali, Zhang et al. 2003, Ran, Bali et al. 2008, Ran, Gadura et al. 2010). These observations underscore the evolutionary versatility of the Hsp90 chaperone as enabler of both robustness and fragility.

However, correlated fitness effects can also be maladaptive. Evidenced by strong genomic signatures, the conserved Hsp90 dependence of maltose and maltotriose metabolism has constrained evolution of maltose and maltotriose metabolism pathways by reducing their sequence space in industrial niches. This is because most *MAL* gene configurations that support efficient traits under basal conditions do not support these traits under conditions of niche-related Hsp90 stress. Hence, Hsp90 both promotes and constrains evolution.

Another important contribution of this study is identification of ecological sources of de-canalization by Hsp90 stress. We show that de-canalization can arise from extrinsic as well as intrinsic Hsp90 stressors. Ethanol is the end product of fermentation and a protein denaturant known to elicit proteotoxic stress (Herskovits, Gadegbeku et al. 1970, Sanchez, Taulien et al. 1992, Alford and Brandman 2018, Voordeckers, Colding et al. 2020) which interferes with beer wort fermentation processes within yeast cells (Carlsen, Degn et al. 1991, Zheng, D’Amore et al. 1994). Ethanol exposure elicits Hsp90 stress at non-toxic industrially relevant concentrations. As a result, ethanol compromises the maturation of Mal63 and reveals cryptic fitness effects of redundant *MAL* gene duplications characteristic of beer and bread yeasts. Notably, the threshold for *MAL* gene copies to canalize maltose metabolism against ethanol stress is higher at increased ethanol concentrations, in line with ethanol being a prime selection agent driving trait canalization in beer and bread yeasts. Added support for this conjecture comes from the substantial amounts of ethanol produced during dough leavening and the interchangeable use of bread and beer yeasts for traditional bread and beer making (Patel and Ingledew 1973, Jayaram, Rezaei et al. 2014). Yet, ethanol is not the only relevant source of Hsp90 stress in this context; heat is also important (Zheng, D’Amore et al. 1994). Relaxed selection for canalization of maltotriose metabolism against Hsp90 stress in lager beer yeasts, as compared to ale beer and bread yeasts, likely reflects the reduced demand on Hsp90 chaperone capacity at the lower temperatures used for lager brewing. These observations suggest that *MAL* gene copy number in beer and bread yeasts is precisely fine-tuned to compensate for the varying severity of niche-related Hsp90 stress. Identifying ecologically relevant Hsp90 stressors and showing how they interact leads us to conclude that Hsp90 stress is a major force of selection shaping the eco-evolutionary dynamics of canalization in nature and demonstrates that context matters in evolution, contrasting the predominant probabilistic-statistical view.

This study has important practical implications. First, the genetic and phenotypic biomarkers of canalization we identified could be useful in strain selection and trait optimization for industrial purposes (e.g., beers with high *vs.* low alcohol content or specific aroma profiles). A notable application of harnessing strains with robust maltose and maltotriose metabolism pertains to biofuel production from sustainable starch-based feedstock in the absence of commercial amylases (Saunders, Izydorczyk et al. 2011). Additionally, principles of robustness uncovered here could be used for the optimization of synthetic traits involving heterologous expression of large multi-meric complexes containing Hsp90 clients (Ignea, Cvetkovic et al. 2011, Denby, Li et al. 2018). As a second application, this work makes predictions about the role of Hsp90 and Hsp90 stress that could be relevant to health and disease. Although caution is warranted when drawing comparisons between budding yeasts and humans, reports indicate that Hsp90 can buffer pathogenic genetic variation in humans and that such buffering can be hyper-sensitive to clinically relevant Hsp90 stressors (Karras, Yi et al. 2017). Our findings suggest that a de-canalization model to explain inter-individual differences in susceptibility to ethanol-related diseases (Sokol, Delaney-Black et al. 2003, Rehm, Gmel et al. 2017). They also suggest a testable hypothesis in which oncogene amplifications can maintain adaptive canalization of oncogenically-activated Hsp90-dependent signaling pathways against proteotoxic stressors often found in the tumor microenvironment (Whitesell and Lindquist 2005, Condelli, Crispo et al. 2019, Lauer and Gresham 2019). Clearly our understanding of the role of canalization in evolutionary processes and human disease is in its infancy (Koonin and Wolf 2009, Fitzgerald, Hastings et al. 2017, Gibson and Lacek 2020, Crespi, Burnap et al. 2022), but our findings highlight the untapped potential of Hsp90-buffered variation as an important source of phantom heritability in ecologically relevant traits.

## Acknowledgments

This work is dedicated to Susan Lindquist who inspired its pursuit. We are grateful to Linda K. Clayton and Luke Whitesell for editing the manuscript, Daniel F. Jarosz, Swathi Arur, Catherine McLellan, Richard Behringer and Pierre McCrea for critical comments on earlier versions of the manuscript. We are also grateful to Ndubuisi Azubuine for technical assistance, Silas Amarell, Wilson Gomarga, Nikitha Kota and Selena Ding for generating yeast growth data, Prathapan Thiru, George Bell, Brant Gracia, Stan Bujnowski, Bo Peng and Jing Wang for bioinformatics support, Tin Chanh Pham for assistance with confocal microscopy and Adriana Paulucci-Holthauzen, core-director for BSRB Microcopy Laboratory, Department of Genetics, who helped with guidance on microscopy acquisition and image processing, Dirk Landgraf and Julia Bonner for providing pDML215 and pDML209 plasmids, Dawn Thompson, Gerry Fink, Duccio Cavalieri and Susan Lindquist for providing yeast strains and antibodies, Joseph Schacherer for protocols on yeast genome purification and scripts for tetrad analysis.

## Author contributions

Conceptualization and Methodology: GIK; Software and resources: DP, HA, WS, NOL, BH; Formal analysis and investigation: DP, HA, YS, NC, BJ, GIK; Validation: DP, HA, GIK; Writing: GIK; Supervision: GIK; Funding acquisition: GIK.

## Data and materials availability

All unique/stable reagents generated in this study are available from the Lead Contact with a completed Materials Transfer Agreement. Further information and requests for resources and reagents should be directed to and will be fulfilled by the lead contact Georgios Karras. All original code is available and will be distributed upon request. CONVICT has been deposited in github (https://github.com/wshropshire/convict).

