## supplemental files for "Ethanol Drives Evolution of Hsp90-Dependent Robustness by Redundancy in Yeast Domestication"

### Materials and Methods

#### Yeast Strain Collection

In this study, 698 yeast strains were phenotyped, 107 of which were sequenced. Strains were obtained from the following sources. 374 natural strains (wild plate 1, wild plate 3, UCD1, UCD2, UCD3), including 204 wine, 21 beer, 8 bread, 9 other fermentation, 65 clinical, 35 soil or fruit, and 32 unknown strains were described previously (Halfmann, Jarosz et al. 2012). 72 *S. cerevisiae* (SGRP Set2) and 83 *S. paradoxus* (SGRP Set 3) strains in their corresponding two mating types and diploid derivatives were described by Liti et al., (Liti, Carter et al. 2009, Bergstrom, Simpson et al. 2014) (purchased from NCYC). 52 beer, 3 whiskey and 8 other fermentation strains were purchased from White Labs. 11 bread and 5 other fermentation strains from maltose-poor fermentation niches were purchased from NCYC. 80 wild strains, including 70 strains from the gut of wasps were a kind gift from Dr. Duccio Cavalieri (Stefanini, Dapporto et al. 2012, Rizzetto, Ifrim et al. 2016). Additionally, 89 derivatives of these isolates were phenotyped, including deletion and fluorescently labelled strains, 36 of which were also sequenced in parallel with their cognate parental strains. 10 isolates from the *Saccharomyces sensu stricto* clade were kindly provided by Dawn Thompson and Gerry Fink. All strains were stored at -80°C in glycerol-containing YPD media (yeast extract 0.67% w/v, peptone 1.33% w/v, glucose 1.33% w/v, glycerol 16.67% v/v).

#### Media and Culture Conditions

Growth and maintenance of the yeast strains were carried on standard YPD media (yeast extract 1% w/v, peptone 2% w/v, glucose 2% w/v) or YPD agar plates. Frozen strains were recovered on YPD agar plates at 30°C and 3–5 single colonies were analyzed. Frozen 96-well yeast glycerol stocks were thawed completely and pinned using 96-pin replicators (Scinomix, Cat# SCI-4010-0S) onto rectangular YPD agar plates containing Ampicillin. To select for strains carrying an antibiotic resistance marker, YPD media was supplemented with 250 μg/mL antibiotics: nourseothricin (JenaBio, Cat# AB-102XL) for marker gene *natMX6*, hygromycin B (Sigma-Aldrich, Cat# H3274) for marker gene *hphMX6*, and geneticin (ThermoFisher, Cat# 10131035) for marker gene *kanMX4*. Metabolic traits were evaluated in YPM (yeast extract 1% w/v, peptone 2% w/v, maltose 2% w/v) or other YP-based media containing the desired carbohydrate in place of glucose. For industrial niche experiments, strains were grown in wort broth (15 g/L malt extract, 0.78 g/L peptic digest of animal tissue, 12.75 g/L maltose, 2.75 g/L dextrin, 1 g/L dipotassium phosphate, and 1 g/L ammonium chloride) and synthetic sourdough media (24 g wheat peptone Sigma-Aldrich, 0.2 g $MgSO_4 \cdot 7H_2O$, 50 mg $MnSO_4 \cdot H_2O$, 4 g $K_2HPO_4$, 1 mL Tween 80, pH adjusted to 4.5 with citric acid) (Bigey, Segond et al. 2021). Sugars and vitamins were added to autoclaved synthetic sourdough media after filter sterilization at 1.5% w/v glucose, 3.5% w/v maltose, and 1 x vitamins (for 1000 X stock: cobalamine, 0.2 g/L; folic acid, 0.2 g/L; nicotinamide, 0.2 g/L; pantothenic acid, 0.2 g/L;

pyridoxal-phosphate, 0.2 g/L and thiamine, 0.2 g/L). All procedures were performed at 30°C unless otherwise indicated.

High-Throughput Phenotyping and Growth Curve Analysis

Metabolic trait efficiency was evaluated as a function of growth in rich media containing a specified carbohydrate in high throughput, similarly to previous work (Warringer, Zorgo et al. 2011). Yeast strains were grown for 24 hours at 30°C in 96-well transparent round-bottom polystyrene microplates (Greiner Bio-One, Cat# 650185) containing 200 μL YPD, and were subsequently diluted (1:50) in duplicate in 384-well clear flat-bottom polystyrene microplates (Corning Life, Cat# 3680) containing 50 μL of YP supplemented with 2% of the indicated carbon source and chemical supplement in each well. Inoculated wells were subsequently mixed using a 384-pin replicator (Scinomix, Cat# SCI-6010-0S), the plates were sealed with clear PCR film (Eppendorf) to eliminate edge-effects due to evaporation and were allowed to grow without plate shaking at 25°C, unless otherwise indicated, in an automated spectrophotometer (Multiskan Sky, ThermoFisher Scientific). The optical density of each culture at 600 nm was measured every 15 minutes over a period of 48 hours, at which stage the experiments were terminated. Experiments with ethanol were setup such that every well in the plate contained the same concentration of ethanol to eliminate cross-well effects of ethanol evaporation over prolonged incubation times. Cells were grown in rich media (YP without selection, unless otherwise noted. The same conclusions were reproduced in synthetic media (CSM), albeit overall yield was lower due to the limiting nutrient conditions. Every experiment included a panel of 6–8 strains that were used as a reference for Hsp90 and ethanol treatment and sugar utilization.
To generate growth curves and extract growth parameters (growth rate, integral, lagging time) we used the growfit-1.1.1 library (Kahm, Hasenbrink et al. 2010) in R (v3.6), and Perl to run custom scripts (available upon request). We removed the first 8 readings from each growth curve to eliminate artifacts originating from bubbles in unsettled cultures and subtracted 0.19 from each $OD_{600}$ reading to account for the starting $OD_{600}$ of the culture. Growth parameters most representative of fitness were strongly correlated with each other, especially the area under the curve (AUC) and maximum growth rate ($R^2 = 0.823$, $p < 0.0001$). As a proxy for metabolic trait efficiency, we utilized quantitative fitness parameters derived from growth curves. Raw data were normalized to positive controls to obtain an estimate of relative growth in each condition. We subsequently normalized these values over the relative growth in YPD to obtain an estimate for metabolic efficiency. For Hsp90i- and ethanol-treated samples, DMSO control and untreated controls were used for normalization, respectively. DMSO had no effect on growth at any temperature. AUC encompasses additional information about growth kinetics not reflected on the growth rate (lag phase duration and maximum yield) (Bell 2010, Sprouffske and Wagner 2016, Ram, Dellus-Gur et al. 2019), thus we chose it as a proxy for metabolic efficiency. To ensure comparability of data across different plates, every experiment was subjected to a stringent technical reproducibility cutoff ($R^2 > 0.98$), including reproducibility across a panel of 6 – 8 strains with various degrees of metabolic robustness to Hsp90i or ethanol (-0.7— -0.1) which were present in each plate, and the relative positions of each strain within the plate was routinely changed to exclude any plate position effects. Every experiment met these reproducibility requirements. All our conclusions were recapitulated also when growth rate was used in place of AUC as a proxy for metabolic efficiency.

Ploidy Estimation

Single colonies were inoculated overnight and grown to an $OD_{600}$ of 0.5 – 0.7 the next day. Cells from 1 mL of each log–phase culture were pelleted by centrifugation at 500 x g at 4°C for 5 minutes, supernatants were removed by aspiration, and cell pellets were washed with 5 mL of sterile water, vortexed, and centrifuges again as above. Cells were subsequently resuspended in ~ 100 μL water and fixed by dropwise addition of 2.5 mL of 100% ethanol and the samples were stored at 4°C overnight. The next day, cells were first washed in 5 mL of 50 mM Tris HCl [pH 7.5] and subsequently were treated with RNase A (380 μg/mL, Invitrogen) in the presence of $MgCl_2$ at 37°C, shaking at 210 RPM (Innova 4430, New Brunswick), for 4 hours. Samples were then washed and treated with Proteinase K (90 μg/mL, Invitrogen) in 50 mM Tris HCl [pH 7.5], incubated at 50°C for 30 minutes. Samples were centrifuged and sonicated (20% power, 3 cycles, 1 second per cycle, Fisherbrand). DNA staining was performed in 96-well plates in 200 μL of 50 mM Tris HCl [pH 7.5] supplemented with SYTOX green (1 μM, Invitrogen). *S. cerevisiae* strains ATCC 200912 (diploid) and S0003 (haploid) were used as references for DNA content. Data were collected using a Guava-5HT flow cytometer (Millipore) and analyzed using FlowJo (v10.8.0).

<u>Yeast Crosses and Tetrad Analysis</u>
Crosses between wild-derived haploid strains (SGRP Set2) and laboratory strain YLK1879 were performed using single colonies of each parental strain on YPD plates and incubated at 30ºC for 24h. Diploids identified by sequential streaking and replica-plating of single colonies onto appropriate selection plates (SD-URA, YPD + hygromycin B). 3–5 single-colonies were evaluated for ploidy by cell cytometry to confirm they are diploid. Diploid strains were grown to saturation in YPD at 30°C and 0.5 mL of saturated culture was washed 5 times in sterile water and incubated in potassium acetate (2%, w/v) for 3–14 days. An aliquot of the sporulated cells was treated with Zymolase (100T, Amsbio) and tetrads were dissected with a micromanipulator (MSM 400 microscope platform, Singer Instruments). Sporulation efficiency was calculated as the ratio between the number of sporulated cells (with 4 spores) in several random sights and the total number of cells in the same sights. Spore viability was tested by dissecting at least 54 tetrads (216 spores) per isolates. A minimum of 8 tetrads were dissected and phenotyped from each cross. Mendelian trait inheritance was concluded for crosses in which at least 70% of all dissected tetrads followed a 2:2 segregation pattern (Hou, Sigwalt et al. 2016).

<u>DNA Amplification, Purification, and Cloning</u>
All plasmids were cloned and purified in the *E. coli* DH5α strain, except for *ccdB* containing Gateway destination vectors, for which the DB3.1 strain was used. For plasmid DNA propagation and cloning, 10–30 μL of cells were mixed with 0.5 μL of plasmid DNA and incubated on ice for 1 minute. Cells were then heat shocked at 42°C for 45 seconds and then on ice for 1 minute, and subsequently resuspended in 100 μL of SOC Media for recovery at 37°C for 30 minutes. Recovered cells were then plated on LB plate supplemented with antibiotics, and the plates were incubated at 37°C for 14 – 18 hours. Plasmids were isolated from bacteria using a PureYield Plasmid Mini Prep System (Promega). For colony PCR on yeast strains, single colonies were picked and resuspended in 20 μL 0.1 mg/mL Zymolase (100T, Amsbio), and after incubation at 37°C for 20 minutes, samples were centrifuges at 800 x g for 1 minute and used for PCR using Taq polymerase (NEB). For DNA cloning and generation of gene disruption cassettes we employed a stitch PCR method using the Phusion High-Fidelity DNA polymerase (NEB). Restriction Digestion and DNA Ligation reactions were performed in 30 μL reactions according to specifications (NEB). In-Fusion Cloning (Takara) and Gateway LR Clonase (Life Technologies) enzyme mixes were used according to manufacturer's specifications. DNA was separated on 1% agarose gels and visualized under a ChemiDoc (BioRad) using RedSafe Nucleic Acid Staining Solution (FroggaBio). DNA extraction from gels

and PCR cleanup were performed using Nucleospin Gel and PCR Cleanup kit (Takara). DNA concentration of purified DNA was measured using NanoDrop at 260 nm. Inserts were validated using restriction digestion (NEB) and Sanger sequencing.
Plasmids pAG25 and pAG32 were a gift from John McCusker (Addgene plasmids #35121 & # 35122; http://n2t.net/addgene:35121 ; RRID:Addgene_35121 & http://n2t.net/addgene:35122 ; RRID:Addgene_35122) (Goldstein and McCusker 1999). Plasmid pJZC522 which was a gift from Wendell Lim & Stanley Qi (Addgene plasmid # 62279; http://n2t.net/addgene:62279 ; RRID:Addgene_62279) (Zalatan, Lee et al. 2015). msfGFP and mRuby3 sequences were described previously (Pedelacq, Cabantous et al. 2006, Bajar, Wang et al. 2016). DNA fragments encoding fluorescent markers were synthesized (IDT) and cloned into plasmid pUC57 pUC57 (Gent), which was a gift from Martin Parniske (Addgene plasmid # 54338 ; http://n2t.net/addgene:54338 ; RRID:Addgene_54338) (Binder, Lambert et al. 2014).

Yeast Genomic DNA Purification

Strains were grown to saturation in 5 mL YPD on a wheel at 25°C and were subsequently collected by centrifugation at 5,000 x g at 4°C for 5 min. All subsequent buffers were prepared using UltraPure water (Thermofisher). Cells were washed once in 1.5 mL of TE [pH 8.0] in 2 mL Eppendorf tubes and the dry pellets were frozen in liquid nitrogen and stored at -80°C. Cell pellets were resuspended in 1,704 $\mu$L Buffer Y1 (1 M Sorbitol, 100 mM EDTA [pH 8.0], Qiagen) supplemented with $\beta$-mercaptoethanol and 2.3 mg Zymolase (100T, Amsbio), and were subsequently incubated at 37 °C for 1.5 hours to generate spheroplasts. Spheroplasts were centrifuged at 5,000 x g at 4°C for 10 min. The supernatant was removed, and pellets were resuspended in 15 mL falcon tubes with Buffer G2 (800 mM guanidine hydrochloride, 30 mM Tris HCl [pH 8.0], 30 mM EDTA, 5% Tween 20, 0.5% Triton X-100, Qiagen) supplemented with 200 $\mu$g RNAseA (Invitrogen) and 400 $\mu$g Proteinase K (Invitrogen). The mixtures were incubated at 50°C for 1 hour by inverting the tubes twice every 10 min. Later, the mixtures were centrifuged at 5,000 x g at 4°C for 10 min. Later 100/G tips (Qiagen) were equilibrated by adding 5 mL of buffer QBT (750mM NaCl, 50 mM MOPS [pH 7.0], 15% isopropanol (v/v), 0.15 % Triton X-100 (v/v), Qiagen). Genomic DNA was bound onto a 100/G column and was washed twice by passing 7.5 mL of buffer QC (1.0 M NaCl, 50 mM MOPS [pH 7.0], 15% isopropanol (v/v), Qiagen) through the column by gravity-flow. Genomic DNA was eluted from the column using prewarmed Buffer QF (1.25 M NaCl, 50 mM Tris HCl [pH 8.5], 15% isopropanol (v/v), Qiagen) directly into 50 mL falcon tubes containing 4 mL isopropanol and 800 $\mu$L sodium acetate (0.3 M). DNA was incubated on ice for 5 minutes and centrifuged at 10,000 x g at 4°C for 20 min. DNA pellets were washed with 1 mL of 70% ice-cold ethanol in 1.5 mL Eppendorf tubes and were subsequently centrifuged at 10,000 x g at 4°C for 10 minutes. Ethanol was removed without disturbing the pellets, and the ethanol wash was repeated twice, after which residual ethanol was completely removed by vacuum centrifugation at room temperature for 1 min. Genomic DNA was resuspended in 50 $\mu$L TE [pH 8.0] and DNA concentration was measure by NanoDrop ($A_{260}/A_{280}$ = 1.8 – 2.2) and the Qubit dsDNA broad range assay kit using a Qubit 4 Fluorometer (ThermoFisher Scientific). DNA quality was evaluated by fragment analysis Bioanalyzer. DNA samples were stored at -80°C.

Library Prep, Whole Genome Sequencing and Genome Assembly

In total 107 strains were sequenced, including a set of 38 engineered derivatives. A sentinel set of strains were sequenced identified 2 or 3 times independently, producing indistinguishable CNV estimates each time. Truseq PCR-free DNA library preparation kit (Illumina) was used to prepare sequencing libraries for each genomic DNA. Prepared libraries were sequenced with

150 nt paired-end sequencing on HiSeqX sequencer (Illumina), producing ~350M PE reads (~105 GB), single index, per lane. Genome assemblies were generated by SPAdes-3.15.5 using the —isolate parameter (median N50 ~350 kb).

Quality Control of raw fastq files and data preprocessing

The FASTQC (v0.11.8) toolkit was performed for quality control of FASTQ files generated from whole genome sequencing (WGS) (Andrews 2010). Clumpify (bbtools v38.33) (Gaia, de Sa et al. 2019) and fastp (v0.23.0) (Chen, Zhou et al. 2018) tools were utilized to remove adapters, low-quality reads and PCR duplicates. Illumina sequencing was performed with 150 bp-long paired-end reads at >100-fold coverage in a single lane. Barcodes were deconvoluted and sequences were mapped on S288C reference genomes. Genome assembly was performed with S288C R64-1-1 (GCA_000146045.2) reference genome (Ensembl) (Cunningham, Allen et al. 2022).

Reference-Based Alignments and Variant Calling

For DNA sequence construction and alignments, we utilized Mac Terminal, Python3, minimap2 (v2.17)– to align reads to a reference genome, Integrative Genomics Viewer (IGV) – to look at alignments, and Flye – to assemble sequenced genome into contigs. For data base searches, sequence and literature, we utilized *Saccharomyces* Genome Database (http://www.yeastgenome.org/) and National Center for Biotechnology Information (http://www.ncbi.nlm.nih.gov/) (Sayers, Bolton et al. 2022). For DNA sequence analysis, creating primers and enzymes, we utilized SnapGene (v5.2.5).

Genetic Distance Based Inferred Neighbor-Joining Tree Analysis

An estimate of genetic distance between the genomes of *S. cerevisiae* (n = 26), *S. paradoxus* (n = 26), and *S. sensu stricto* (n = 10) were calculated using Mash-v2.3 (Ondov, Treangen et al. 2016) and/or FastANI-v1.32 (Jain, Rodriguez et al. 2018). Except for *S. sensu stricto* (Mash distance only), distance matrices produced from mash and fastani were combined using the combine_distance_matrices.py script (Wick, R. Bacsort). These distance matrices were used as inputs to create a neighbor joining tree using the BIONJ algorithm (Gascuel 1997) from the ape-v5.6-1 R package (Paradis and Schliep 2019). Dendrograms were created using the ggtree-v3.3.1 R package (Yu 2020).

Yeast Genome Engineering

To target *MAL* genes, we avoided double-strand break-mediated genome engineering methods such as CRISPR/Cas9-based editing: we reasoned that introducing DNA breaks into highly similar genes, which are sub-telomeric and in close linkage with other non-essential genes (Brown, Murray et al. 2010), would likely promote recombination between the highly similar broken *MAL* gene copies. Such recombination would provoke the formation of off-target structural variations, such as unbalanced reciprocal translocations (Fleiss, O'Donnell et al. 2019, Agier, Fleiss et al. 2021). Hence, to disrupt copies of *MAL* genes while minimizing the probability of unwanted alterations we turned to classical genome editing involving homology-based integration of dominant selection markers (Wach 1996). Although more time-consuming, this approach is less likely to induce translocations or other mutations because it relies on spontaneous single strand breaks or gaps that occur naturally during DNA replication. In addition, this approach targets one gene at a time, rather than not multiple genes thus, allowing functional evaluation of individual gene copies. Similar approaches have been successfully utilized to engineer polyploid industrial or wild strains with efficiencies comparable to CRISPR/Cas9-based strategies (Wach 1996, Generoso, Gottardi et al. 2016).

We employed 450-850 bp-long homology arms (HA-L and HA-R) mapping to the 5' and 3' ends of the target ORF (Figure S3O), amplified by PCR using BY4741 strain as templates. PCR fragments containing an antibiotic selection cassette (NAT: *natMX6*, HYG: *hphMX6*, KAN: *kanMX4*) were generated. All PCR products gel purified and were subjected to a second round of PCR to stitch 2 HA fragments and a selection marker together, using previously described S1 and S2 sequences as overhangs (Longtine, McKenzie et al. 1998, Knop, Siegers et al. 1999, Janke, Magiera et al. 2004). For *MAL63* disruptions in industrial strains, *MAL63* amplicons from the corresponding parental strain were used as templates for stitch PCR. BY4741 was also used as template for all *MAL11* disruptions. For *MAL31* and *MAL32* (almost identical to *MAL12*), we used chromosome II of the BY4741 for most strains, except for the strains used for the baking lineage for which we used genomic DNA extracted from the corresponding parent to generate specific homology arms, as for *MAL63*. PCR reactions were confirmed at each step by agarose gel electrophoresis. First step PCR fragments were gel purified and cleaned before subsequently stitching them together. Generated gene disruption cassettes were precipitated by sodium acetate (0.3 M) and ethanol (80%, v/v) at -20°C for 30 minutes and centrifugation at 16,000 x g at 4°C. Supernatant was removed by aspiration and the DNA pellet was dried by vacuum centrifugation at room temperature for 4 minutes (SpeedVac Savant DNA120, Thermo Scientific). DNA was then resuspended in 20 μL of TE buffer and transformed into competent yeast cells. Deletions were validated by colony PCR and whole-genome sequencing.
Preparation of yeast competent cells and transformation was based on a protocol developed by Gietz and Woods 2002 (Gietz and Woods 2002). Saturated cultures from a single colony were grown to mid-log phase at 30°C, 210 RPM (Innova 4400 incubator shaker, New Brunswick), in 50 mL of YPD. The cells were then collected and centrifuged at 500 x g for 5 minutes at room temperature, washed in sterile water, and SORB buffer (100 mM LiOAc, 10 mM Tris HCl [pH 8.0], 1 mM EDTA, 1 M Sorbitol), and subsequently resuspended in 360 μL of SORB supplemented with 40 μL heat-denatured Salmon Sperm. Competent cells were aliquoted and stored at -80°C. Cell transformations were performed as follows. Competent cells were thawed at room temperature, and 50 μL of the yeast cells were used, along with 10 μL of digested plasmid or stitched PCR product. 360 μL of PEG buffer (100 mM LiOAc, 10 mM Tris HCl [pH 8.0], 1 mM EDTA, 40% PEG 4000) was added to each transformation, vortexed gently, and they were incubated at room temperature for 30 minutes. After the incubation, 46.67 μL of DMSO was added to each transformation, vortex briefly, and incubated at 42°C for 5 minutes exactly. Transformations were spun down at 800 x g in a microcentrifuge for 3 minutes and supernatant was carefully removed. For auxotroph selections, the transformations were resuspended in 200 μL of water and plated on the selected dropout media plate. For antibiotic selections, the transformations were resuspended in 700 μL of YPD and incubated at 30°C, 800 RPM, in a benchtop thermomixer for 3 hours. After incubation, cells were spun at 800 x g for 3 minutes, supernatant was removed, cells were resuspended in 200 μL of water and then plated on YPD-antibiotic plates. For sequential disruption of gene copies, selected colonies were sequentially transformed with different selection cassettes stitched to the same homology arms, at each stage selecting for each cassette that had already been selected by replica plating onto the corresponding antibiotic-containing plates. This was performed to exclude strains in which the disruption cassette had re-integrated into already disrupted alleles.

<u>Total Protein Precipitation from Yeast Cell Extracts</u>
While tube is sitting in ice, add 1000 μL of prechilled distilled water. Add 150 μL of 1.85 N Sodium Hydroxide (Sigma-Aldrich)/7.5% β-mercaptoethanol (Sigma-Aldrich) solution to each sample and briefly vortex. Incubate sample for 15 minutes prior to adding 150 μL of 55% (w/v)

Trichloroacetic acid (Sigma-Aldrich) and briefly vortex. Incubate sample for 10 minutes. Spin samples at 12000 G for 10 minutes. Remove supernatant and do a second spin for 10 minutes. Remove remaining supernatant before adding 50 µL of loading dye (200 mM Tris HCl [pH 6.8], 8 M Urea, 5% SDS, 10 mM β-mercaptoethanol) to each sample and incubate for 10 minutes shaking on a thermomixer (Eppendorf) at 1,400 RPM for 10 minutes. Spin samples at max speed for 15 seconds prior to loading into polyacrylamide gel (Invitrogen).

Polyacrylamide Gel Electrophoresis and Western Blotting

All samples were run in 4-12% 20-well polyacrylamide gel in gel running system (BioRad) filled with 1x NuPAGE running buffer (Sigma-Alrich) for 1 hour at room temperature. The gel was transferred after a brief wash with distilled water onto a PVDF transfer stack (ThermoFisher). The transfer was performed in the iBlot transfer system for 7 minutes. Wash newly transferred membrane in PBS-Tween 20 solution for 5 minutes. Membraned were blocked in PBS-Tween 20, 5% milk for 1 hour and subsequently membranes were incubated with primary antibodies (1:2,000 for anti-GFP antibody Living Colors, 1:5,000 for anti-yHsp90 antibody, 1:50,000 for anti-yPgk1) overnight at 4°C. Membranes were washed in PBS-Tween 20 and incubated with secondary antibody (used at 1:5,000 dilution) in PBS-Tween 20, 5% milk for 45 minutes. Washed in PBS-Tween 20 and developed using the ECL reagent under a ChemiDoc (BioRad).

Confocal Microscopy

Strains expressing fluorescent proteins were grown in minimal media (CSM + sugar) to mid-logarithmic phase and washed in PBS. Diluted cells were placed on slides (Shandon™ ColorFrost™ Plus Slides) and then covered with coverslips (12 mm, Millipore Sigma A-003-E) that had been pre-coated with poly-D-lysine (Millipore Sigma A-003-E). Excess liquid was removed with a Kimwipe, and slides were sealed with nail polish to avoid evaporation. Sealed slides were kept in an inverted orientation for at least 10 minutes to allow for yeast cells to adhere onto the treated coverslip. Confocal images were acquired with a Nikon A1 confocal using a 60X, 1.4NA plan-apo oil objective and a pixel size of 0.2 um.  We used 488nm laser line combined with 525/50nm emission filter for "green probe" (EGFP for $^{GFP}$Mal63, msfGFP for lineage tracing in competition assays) and 561nm laser line combined with 595/50nm for "red probe" (mRuby3). Z-stack acquisition was performed with z-steps of 0.3 uM, while the pinhole was set to 1 Airy unit.  3D rendering was performed with NIS-Nikon Elements.

Competitive Fitness Assays

To evaluate relative fitness, we employed cell competition assays, as they report on differences in the rates of growth and mortality between strains at lag, logarithmic and stationary phases of the culture. To track competing strains, we stably expressed fluorescent proteins (“green”, msfGFP, vs. “red”, mRuby3) under the strong constitutive promoter (GPD). Expression of fluorescent proteins was confirmed by cell cytometry using an Attune NxT Accoustic Focusing Cytometer (Thermo) or a FACSCantoII (BD), with appropriate laser and filter settings for detection of msfGFP (laser 488 nm excitation, detector BL1 channel B530 530/30) and mRuby3 proteins (laser 561 nm excitation, detector YL1 channel YG585 585/16). At least 3 independent isolates of each labeled strain were validated, all of which behaved indistinguishably from their cognate parent strains in growth assays. The relative fitness of competing strains was determined in pairwise or multi-strain competition experiments in mixed cultures at 25°C, as follows.

Strains were inoculated from single colonies and were individually grown for 18-24 hours. For pairwise competition assays, aliquots from saturated cultures of 2 labeled strains were mixed in ratios of 1:1 (pairwise red vs. green, inoculated at $OD_{600}$ 0.02, Figure S5D) or 1:999 (pairwise red vs. green, inoculated at $OD_{600}$ 0.02, Figures S5E and S5F). Multi-strain competition assays involved an unlabeled strain in addition to the two fluorescently labeled strains, which were mixed in 1:1:98 ratio (red vs. green vs. unlabeled, inoculated at 5 x $10^6$ cells per mL, ~$OD_{600}$ 0.3, Figures 5D-I and S5G-L). Mixed cultures were grown to saturation for 40 – 96 hours at 25°C (~ 5.4–10.5 generations) in YPM, wort, or sourdough media supplemented with Hsp90 inhibitor, ethanol, or vehicle control, as indicated. Next, aliquots set to a final $OD_{600}$ ~ of 0.5 in PBS were analyzed by cell cytometry, as described above. A minimum of 10,000 cells were counted from each sample. Gating parameters were optimized based on individual cultures of unlabeled and labeled strains, under basal conditions as well as under the conditions of Hsp90 stress employed in each corresponding experiment. Subsequently, saturated cultures were diluted in fresh media ($OD_{600}$ 0.02, or 0.3, depending on the experimental setup described above), and were allowed to grow to saturation and analyzed by cell cytometry, as above. Experiments were terminated after 3 – 7 rounds of serial dilutions (up to > 50 generations).

Statistical Analyses

Standard statistical analyses were conducted in GraphPad Prism (v9.1) or R studio (2022.07.0+548) with custom scripts. For Hypergeometric analysis we created a custom script that utilizes the hypergeo function in R Studio. P values below 0.05 were considered statistically significant. GraphPad Prism was used for data visualization.

Calculation of Robustness Estimates

Robustness of a metabolic trait to Hsp90 stress, ethanol or genetic variation was defined as the $\log_{10}$ ratio of metabolic efficiency in the presence of the perturbation over the basal efficiency. Metabolic efficiency was calculated as follows. AUC values of target strain-conditions (AUC target) were first scaled relatively to the metabolic efficiency of control strains in the same condition (AUC control) ("relative growth", 0-1 range), using equation (1):

$$\text{rel. growth} = \frac{\text{AUC target}}{\text{AUC control}} \; (1)$$

Relative growth values for each other condition were normalized to the corresponding relative growth values in glucose to obtain metabolic efficiency estimates, using equation (2):

$$\text{Efficiency} = \frac{\text{AUC target}}{\text{AUC control}} \; (2)$$

A strain-condition was deemed positive of growth, and thus the corresponding strain capable of utilizing the provided sugar, if both relative growth and efficiency estimates were greater than 0.25. Robustness estimates (*R*) were calculated only for these strain-conditions with above threshold growth as described for the robustness of maltose utilization to Hsp90 inhibition, using equation (3):

$$R = \log\left(\frac{\text{rel. growth in maltose, Hsp90i}}{\text{rel. growth in maltose, DMSO}}\right) - \log\left(\frac{rel.\, growth\ in\ glucose, Hsp90i}{rel.\, growth\ in\ glucose, DMSO}\right) \; (3)$$

where "rel. growth in maltose, DMSO" and "rel. growth in glucose, DMSO" > 0.25.

Clustering Methodologies

Clustering was performed using Cluster 3.0 (de Hoon, Imoto et al. 2004), using a centered similarity metric and centroid linkage clustering method. Dendrograms were generated with TreeView (v1.16r4) (Saldanha 2004) and heat maps using GraphPad Prism (Figure 3A). Strains in Figures S3E and S3F were parsed into two groups based on the robustness of maltose metabolism against moderate-level Hsp90 inhibition (radicicol, 10 μM) using a robustness cutoff of -0.2 (kmeans, R).

Calculation of Coverage Maps

To identify deviations in coverage across the genome, reads from each sample were aligned to the reference assembly using bwa and the resultant bam file was sorted with Samtools (v1.14) (Li, Handsaker et al. 2009, Danecek, Bonfield et al. 2021). Chromosomal coverage was split into bins of 1,000 bp using bbmap pileup.sh (Bushnell, Rood et al. 2017). Subsequently, the reads were aligned to the control gene set with bwa, the resultant bam file was sorted with Samtools, and the coverage was calculated using bbmap pileup.sh. To find the average baseline coverage of the genome, agnostic of copy number events, the coverage of the control genes was averaged. Each chromosomal coverage bin was divided by the average control gene coverage to get a copy number ratio representing each bin. Plots were made by plotting position on the x axis (chromosome number) and normalized coverage on the y-axis.

Estimating Gene Copy Number Variations

Two independent tools were utilized to estimate variations in gene copy number across different genomes: 1) Control-FREEC (v11.6) (Boeva, Popova et al. 2012) calculates coverage by dividing the genome into fixed non-overlapping windows (i.e., 100 bp) using a LASSO-based algorithm and counting reads within each window, applying LOESS modeling for GC-bias correction. Implemented Control-FREEC parameters included window = 100, minExpectedGC = 0.35, maxExpectedGC = 0.55, telocentromeric = 7,000 and breakPointThreshold = 0.8. The Control-FREEC results were annotated with Samtools (version 1.14). Mean of ratios of all bins for ORFs are calculated in RStudio (version 4.0.3) ((2020) 2020). 2) COpy Number Variant quantifICation Tool (CONVICT) is described in this study (CONVICT-v1.0 is available on github: Shropshire W, CONVICT: GitHub: https://github.com/wshropshire/convict). CONVICT queries a target ORF database of interest, segments each ORF into fixed non-overlapping bins (i.e., 100 bp), and calculates the ratio of coverage depths between target and control ORFs. From these bins we calculated a coefficient of variation (CV) defined as (standard deviation of bin coverage)/(mean bin coverage) to measure the binned coverage dispersion in order to control for sequencing artifacts. Bins with coverage above the mean coverage plus or minus one standard deviation were removed and this was done iteratively until the gene's CV reached a user defined threshold. The CV threshold was determined by selecting a set of 20 target and 20 control genes with known copy number deviations in control genomes and conducted the analysis at CV threshold increments of 2.5% starting from 0% to 25%. We found that at a level of 17% artifacts were removed while maintaining real biological signal. The primary output is an estimate of normalized coverage depths, or relative copy number. Absolute copy number estimates are calculated by multiplying the relative copy number of each gene or ORF derived from either tool to the ploidy of the corresponding strain, as determined by cell cytometry. The X12576.1 sequence was used as reference for *S. pastorianus MAL63.*

We applied multiple configurations of each tool corresponding to different read-normalization methodologies and accounting for GC content, mappability, use of different control genomes with well characterized gene copy numbers, and different definitions of control genes. We used various criteria to identify control genes in the nuclear genome, 1) essential housekeeping genes located away from centromeres and telomeres, 2) no known pseudogenes, 3) genes with significant gene dosage effects assuming that variability in copy number of these genes reduces fitness across different genomic backgrounds (Burke, Gasdaska et al. 1989, Makanae, Kintaka et al. 2013), 4) 651 least variable genes in copy number our dataset across based on principal component analysis. Average coverage of control genes was proportional to strain ploidy. We observed very similar results across all different approaches. We avoided the use of neighboring genes as controls (Bigey, Segond et al. 2021) because many are in linkage disequilibrium with *MAL* genes. Our conclusions were robust to any combination of these permutations and were also recapitulated using both CNV analysis tools.

High similarity between different *MAL* genes in the same family can introduce noise during allocation of reads to different genes which can affect the accuracy of copy number estimation. To address this issue, we incorporated a clustering step into CONVICT. Similar ORFs were clustered based on usearch (v11.0.667) (Edgar 2010) to consolidate highly similar ORFs into consensus target gene families based on a 90% identity threshold. CONVICT generates a list with the identity of the target genes clustered together into a gene family ("C" in column A indicates separate clusters and numbers identify the original target gene list, "clusters.tsv"). CONVICT aligns raw reads to the clustered target genes and control gene sequences using bowtie2 aligner (v2.4.1) (Langmead and Salzberg 2012) before segmenting ORFs into non-overlapping bins. Mapping reads from the almost identical paralogues, *MAL32* and *MAL12*, onto a gene family consensus sequence (*MALx2*) eliminated outliers and drastically increased the fraction of absolute copy number estimates approximating integer numbers. We found this clustering-based approach was more accurate than summing up the un-clustered data although the latter outperformed CNV estimations neglecting to account for sequence similarity. Copy numbers of *MAL* genes in disrupted lineages were estimated for each strain by averaging the coverage of central 100-bp windows within the deleted regions within the ORF and then normalizing these values with respect to the average coverage of control ORFs. Further normalization of copy number estimates against the average copy number of the parental strains gave the relative copy number estimates plotted in Figures 3D and S3S. Hybrid strains were identified using taxonomic sequence classification systems Kraken 2 (Wood, Lu et al. 2019) and Bracken (Lu, Rincon et al. 2022). All conclusions were robust to exclusion of these strains from downstream analyses.

<u>Adjusting Copy Number Estimates Based on Ploidy</u>

Relative gene copy number was chosen as more representative of the biology of metabolic traits, as compared to absolute gene copy number because it accounts for the effect of ploidy on cell size. Given that cell volume increases proportionately with ploidy in yeast (Mortimer 1958), while protein concentration remains, on average, unchanged (de Godoy, Olsen et al. 2008), it is obvious why the absolute copy number of i.e. metabolic genes overestimates the effects these gene copies evert on metabolic flux and fitness, as compared to the intracellular concentration of the encoded proteins (Keren, Hausser et al. 2016). Hence, relative gene copy number estimates (normalized over control genes reflecting ploidy) were preferred over absolute gene copy number estimates.

Nevertheless, although more physiologically relevant than absolute gene copy number, relative copy number is not perfectly correlated with the concentration of the encoded proteins within the cell, and thus our analysis has a few caveats that must be accounted for. First, it remains unknown if protein concentration is similar across ploidy states beyond the diploid in yeast. Second, membrane proteins increase in amount upon diploidization proportionately with cell surface, not cell volume (Comai 2005, de Godoy, Olsen et al. 2008). Third, heterogeneity in cell size and its determinants beyond ploidy (i.e., cell wall integrity, cell cycle) are likely prevalent in natural populations. Hence, while the concentration of the cytosolic proteins Malx2 and Malx3 within the cell are not expected to differ very much between diploid and tetraploid cells, the concentration of membrane localized Malx1 permeases is expected to be reduced in tetraploids by a factor of ~ 0.79, as compared to diploids, in agreement with geometrical considerations. Nevertheless, the above considerations only emphasize further the importance of *MALx2* and *MAL63* in the canalization of industrial traits in domesticated yeasts.

Gene-Set Enrichment Analysis for Copy Number Variations

Gene-set enrichments between various groupings of yeast strains were determined using relative CONVICT CNV calls for each gene-ontology (GO) term. Custom gene sets (maltose utilization and palatinose utilization pathways) were included in addition to standard GO term definitions (Chen, Tan et al. 2013, Kuleshov, Jones et al. 2016). Gene-set copy numbers were found by summing the gene copy numbers of each set member, producing a gene-set copy number. False discover rate Multiple comparison analyses were performed using 2-tailed independent sample t-tests with a Bonferroni correction to produce the plotted adjusted p-values. The same methodology was used to identify individual gene level differences between strain types. Figure S3E was generated using K-means clustering to group strains into sensitive (N = 19) and robust (N = 39) to Hsp90 stress for maltose utilization and compared the average gene copy number across GO term categories.

Calculation of Selection Coefficients

Selection coefficients (*s*) were calculated as previously described (Hegreness, Shoresh et al. 2006, Chevin 2011), based on the following equation,

$$s = \frac{1}{t}\ln\left(\frac{\text{ratio } tf}{\text{ratio } tf - 1}\right), \qquad \text{ratio} = \frac{\text{frequency of competing strain}}{\text{frequency of outcompeted strain}} \qquad (4)$$

where $t_f$ the examined time-point (~6.6–50 generations) and $t_{f-1}$ is the preceding competition round (Dykhuizen 1990). The selection coefficient was estimated as an average of 4 – 16 independent competition assays (Teresa Avelar, Perfeito et al. 2013). In addition to relative differences in fitness, cell competition assays also report on synergistic or antagonistic interactions between strains (Concepcion-Acevedo, Weiss et al. 2015). No strain-strain interactions were observed.

### Fig. S1. Ecologically relevant traits vary in their robustness to Hsp90 impairment.

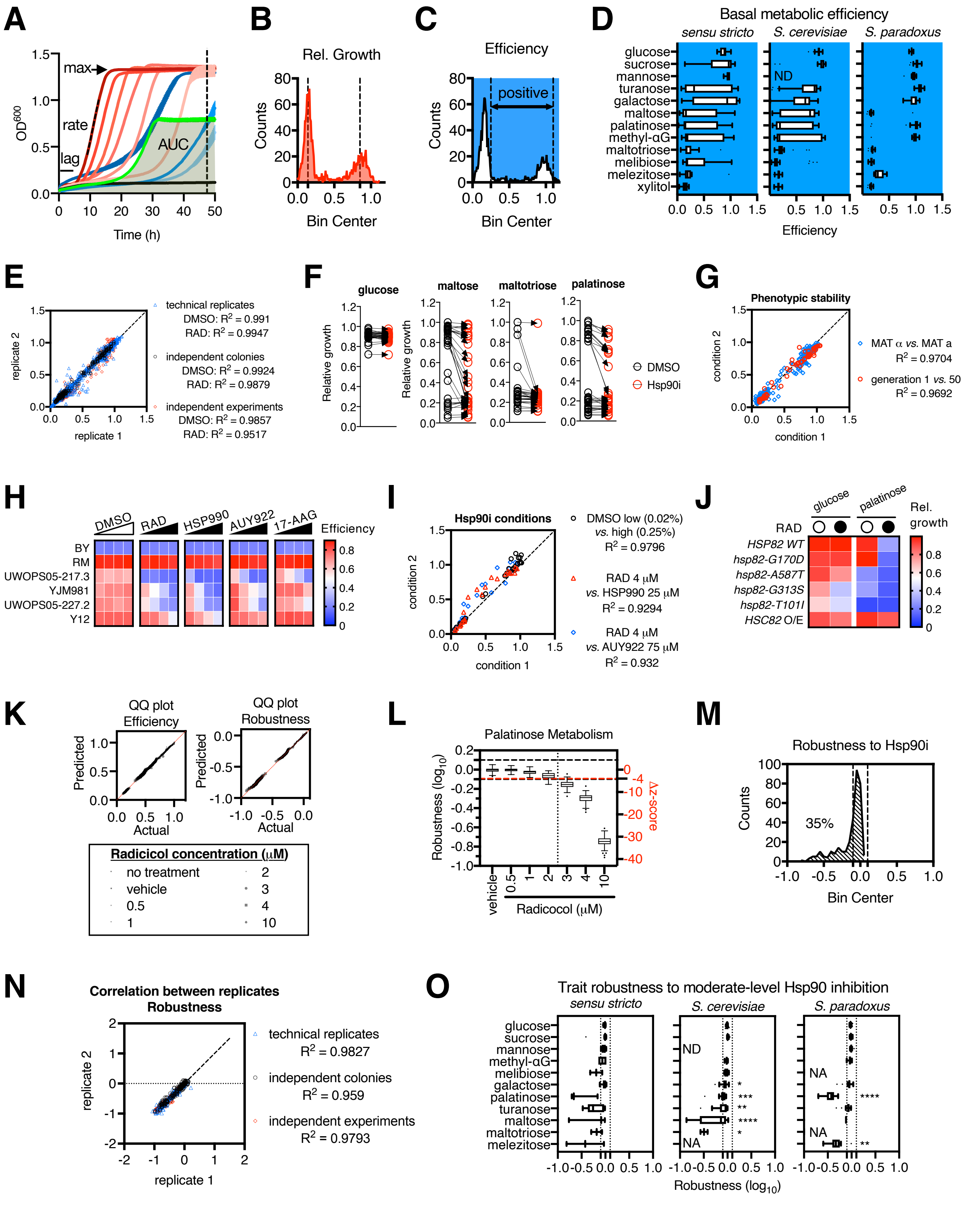

**Figure S1. Ecologically relevant traits vary in their robustness to Hsp90 impairment.** (**A**) Reproducibility between replicate growth curves generated from independently inoculated cultures in maltose media. Three *S. cerevisiae* isolates (red, blue, green) were grown in the presence of different concentrations of radicicol (0.1 – 10 μM, from dark to light color). Growth parameters indicated: lag phase, growth rate, maximum yield, area under the curve (AUC, shaded). Error bars are smaller than the symbol and represent standard deviations from duplicate growth curves. (**B** and **C**) Frequency distribution of (B) relative growth and (C) efficiency estimates for 11 metabolic traits across 170 *Saccharomyces* strains (including isogenic haploid strains of different mating type and diploid derivatives) in the presence of low (4 μM) *versus* moderate radicicol concentrations (10 μM) *versus* the corresponding concentrations of vehicle control (DMSO). Lower threshold (left dotted line) identifies strains with significant metabolic efficiency (> 25%, z > ~3, "positive"). (**D**) Box plots of basal metabolic efficiency of 11 metabolic traits (in addition to glucose utilization) across 10 *sensu stricto,* 26 *S. cerevisiae*, and 23 *S. paradoxus* isolates. Whiskers based on Tukey method. "ND" indicates not determined. (**E**) Correlation between technical replicates (independent cultures in the same plate) and biological replicates (independent colonies and independent experiments). Growth data from strains treated with different radicicol concentrations were pooled (RAD). All correlations are significant. (**F**) Before-after plots of the efficiency of the indicated metabolic traits under basal (DMSO, vehicle control) or Hsp90 inhibition conditions (Hsp90i, radicicol 10 μM). (**G**) Correlation between metabolic efficiency under basal and Hsp90 inhibition conditions determined at different generation numbers of the culture or between different mating types. (**H**) Strain-specific sensitivity of maltose metabolism to small molecule Hsp90 inhibitors from distinct structural groups: resorcinol (radicicol: 1 μM, 2.5 μM, 5 μM, 10 μM, AUY922: 10 μM, 25 μM, 50 μM, 75 μM), isoxazole resorcinol (HSP990: 10 – 75 μM), ansamycin benzoquinone (17-AAG: 10 – 75 μM). (**I**) Correlations between metabolic efficiency values determined in the presence of the indicated small molecular Hsp90 inhibitors. (**J**) Relative growth of laboratory strain ATCC 200912 expressing different Hsp82 variants expressed as sole source of Hsp90 protein under the control of the GPD promoter or wild-type Hsc82 protein under its endogenous promoter. Filled spheres represent administration of radicicol (10 μM). (**K**) QQ-plot for normality of efficiency and robustness to Hsp90 inhibition of palatinose metabolism in laboratory strain ATCC 200912; all conditions passed two (D'Agostino & Pearson and Kolmogorov-Smirnov) or more normality tests. Negative robustness values signify the degree of sensitivity of palatinose metabolism to radicicol treatment. (**L**) Box plots indicate variance of robustness estimates for each of 96 single colonies of strain ATCC 200912 growing in parallel in palatinose media supplemented with the indicated concentrations of radicicol *versus* vehicle control. Vertical dotted line indicates threshold of significant differences in robustness upon radicicol treatment as compared to vehicle control (Δz-score < -4). (**M**) Frequency distribution of robustness to low or moderate radicicol treatment across all metabolic traits and conditions as in panel D. (**N**) Correlation of robustness estimates between technical and biological replicates. (**O**) Box plots of robustness of 11 metabolic traits to radicicol across 10 *sensu stricto,* 26 *S. cerevisiae*, and 23 *S. paradoxus* isolates. All coefficients of determination ($R^2$) are statistically significant (Pearson, $p < 0.0001$). All values represent averages derived from 2 – 3 independent experiments. Significance between distributions was determined using the Kruskal-Wallis test with Dunn's multiple comparisons.

**Fig. S2. Differential robustness of traits against Hsp90 inhibition in domesticated *versus* wild yeasts.**

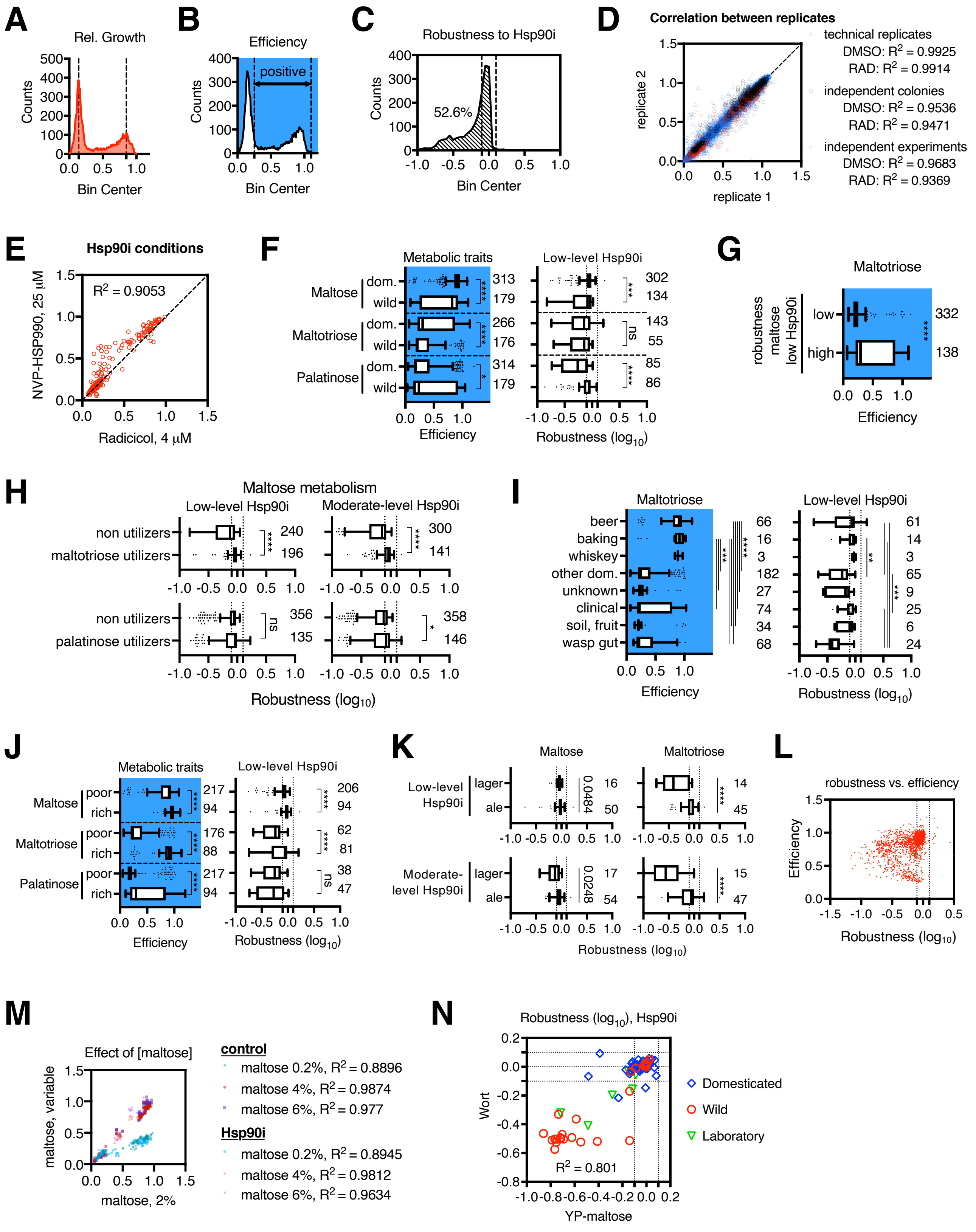

**Figure S2. Differential robustness of traits against Hsp90 inhibition in domesticated *versus* wild yeasts.** (**A-C**) Frequency distributions of (A) relative growth, (B) efficiency, and (C) robustness to radicicol treatment across 534 natural isolates under 4 conditions (radicicol low *vs.* moderate, and corresponding DMSO controls). (**D**) Correlation between technical and biological replicate metabolic efficiency estimates. (**E**) Correlation between maltose utilization efficiency under conditions of Hsp90 inhibition with chemically distinct small molecule Hsp90 inhibitors. (**F**) Box plots of basal efficiency (left panel) and robustness to low-level Hsp90 stress (right panel) for maltose, maltotriose, and palatinose metabolism comparing domesticated (wine, beer, bread, distillation, sake, cider, etc.) and wild (clinical, soil, fruit, wasp gut) strains. Whiskers based on Tukey method. (**G**) Box plot of efficiency of basal maltotriose metabolism across strains parsed by the robustness of maltose utilization to Hsp90 stress. (**H**) Box plots of the robustness of maltose utilization to low- *versus* moderate-level Hsp90 stress parsed by maltotriose or palatinose utilization efficiency. (**I**) Box plots of basal efficiency (left panel) and robustness of maltotriose utilization to Hsp90 stress (right panel) parsed by strain origin. (**J**) Box plots of basal efficiency (left panel) and robustness (right panel) of the indicated metabolic traits to low-level Hsp90 stress for industrial strains parsed by prevalence of maltose in the domestication niche (poor *vs.* rich). (**K**) Box plots of robustness of maltose (left panels) and maltotriose metabolism (right panels) to low-level (radicicol 4 µM, upper panels) and moderate level Hsp90 stress (radicicol 10 µM, lower panels) comparing industrial ale and lager beer strains. Significant differences indicated with asterisks (*). Differences in robustness of maltose metabolism were not significant after correction for multiple hypothesis testing. (**L**) Correlation between basal efficiency and robustness to Hsp90 stress across all metabolic traits and strains in the entire dataset. (**M**) Correlation between maltose utilization efficiency under basal conditions and under low-level Hsp90 stress across different maltose concentrations. Limiting the maltose concentration reduces metabolic efficiency but does not affect robustness to Hsp90 stress. (**N**) Correlation of robustness to low-level Hsp90 stress between maltose and wort utilization across strains of diverse origins. Values represent averages of 2 – 3 independent experiments. All coefficients of determination ($R^2$) are statistically significant (Pearson, $p < 0.0001$). Significance between distributions was determined using Mann-Whitney or Kruskal-Wallis test with Dunn's multiple comparisons.

**Fig. S3. Genetic determinants of trait robustness to gene loss and Hsp90 stress.**

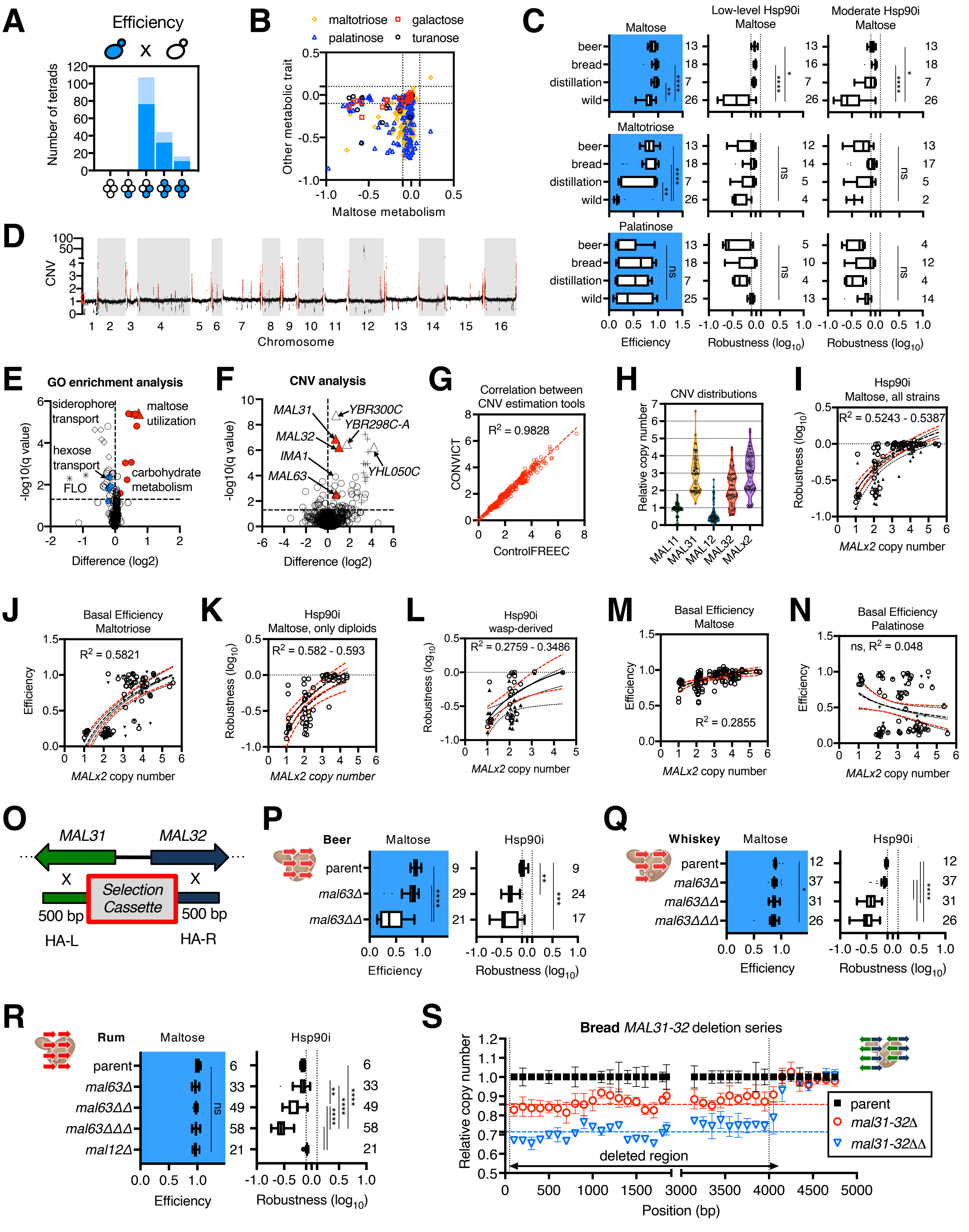

**Figure S3. Genetic determinants of trait robustness to gene loss and Hsp90 stress.** (**A**) Summary of tetrad analysis for 11 crosses between wild-derived haploid parent strains. Blue spores in the tetrad arrangement indicate pattern of inheritance of basal efficiency of maltose metabolism. Light blue bars indicate the fraction of tetrads for which the robustness of this trait differed as compared to the maltose utilizing parent. (**B**) Comparison between robustness of maltose utilization against the robustness of other indicated metabolic traits to moderate-level Hsp90 stress. (**C**) Box plots of basal efficiency and robustness to low- and moderate-level Hsp90 stress for maltose, maltotriose, and palatinose utilization across sequenced strains based on origin. (**D**) Whole-genome coverage maps relative to the haploid S288C reference genome averaged at 1kb bins across 64 natural yeasts. Averages are indicated by black spheres and standard deviations are shown in red; most standard deviations are smaller than the symbol. (**E** and **F**) Volcano plots of CNVs associated with robustness of maltose metabolism to moderate-level Hsp90 stress (radicicol, 10 µM) using a cutoff of -0.2 analyzed at the level of (E) pathways (GO term category average) and (F) individual ORF (CONVICT no cluster). Indicated categories: maltose utilization (red triangle), carbohydrate metabolism (red spheres), flocculation (*FLO1*, asterisks), hexose transport (blue spheres), siderophore transport (diamonds), Ty retrotransposon (crosses). Multiple unpaired t-tests of $Log_2$ transformed data with two-stage step-up multiple correction method FDR 5%. (**G**) Correlation between CNV estimates for *MAL* genes between CONVICT and Control-FREEC. (**H**) Absolute copy number of indicated *MAL* genes as determined by CONVICT without (*MAL12*, *MAL32*) *versus* with clustering (*MALx2)*. Results for *MAL11* and *MAL31* were identical using both methods. (**I-N**) Semilog plot of interpolation fit between *MALx2* relative copy number and (I) robustness of maltose metabolism to moderate-level Hsp90 stress, (J) basal efficiency of maltotriose metabolism, (K and L) robustness of maltose metabolism to moderate-level Hsp90 stress across sequenced (K) diploid strains, *versus* (L) strains derived from the gut of wasps, and basal efficiency of (M) maltose and (N) palatinose metabolism across all sequenced strains. (**O**) Schematic of homology-directed gene disruption using selection cassettes with homology arms flanking the targeted disrupted sequence (left, HA-L or right, HA-R). (**P-R**) Genotype schematic and box plots of basal efficiency and robustness of maltose metabolism to moderate-level Hsp90 stress in *mal63* disrupted lineages derived from industrial strains (P) beer, (Q) whiskey, and (R) rum. (**S**) CNV analysis of *MAL31-MAL32* relative copy in commercial bread strain and derivatives. 7 copies of the *MAL31-MAL32* pair are present in this strain. Disruption of each *MAL31-MAL32* pair is expected to reduce the gene's coverage in the deleted region by ~ 0.1429 (1:7, expected coverage *Δ*: 0.8571, and *ΔΔ*: 0.7143) as compared to the corresponding parental strains. Relative coverage is shown at each non-overlapping 100 bp-windows spanning the length of the ORFs for 2 – 7 replicate strains per genotype. Values and standard deviations were derived from 2 – 3 independent replicate experiments. Significance between distributions was determined using Mann-Whitney or Kruskal-Wallis test with Dunn's multiple comparisons.

**Fig. S4. Niche-related proteotoxic exposures phenocopy low-level Hsp90 inhibition in perturbing traits.**

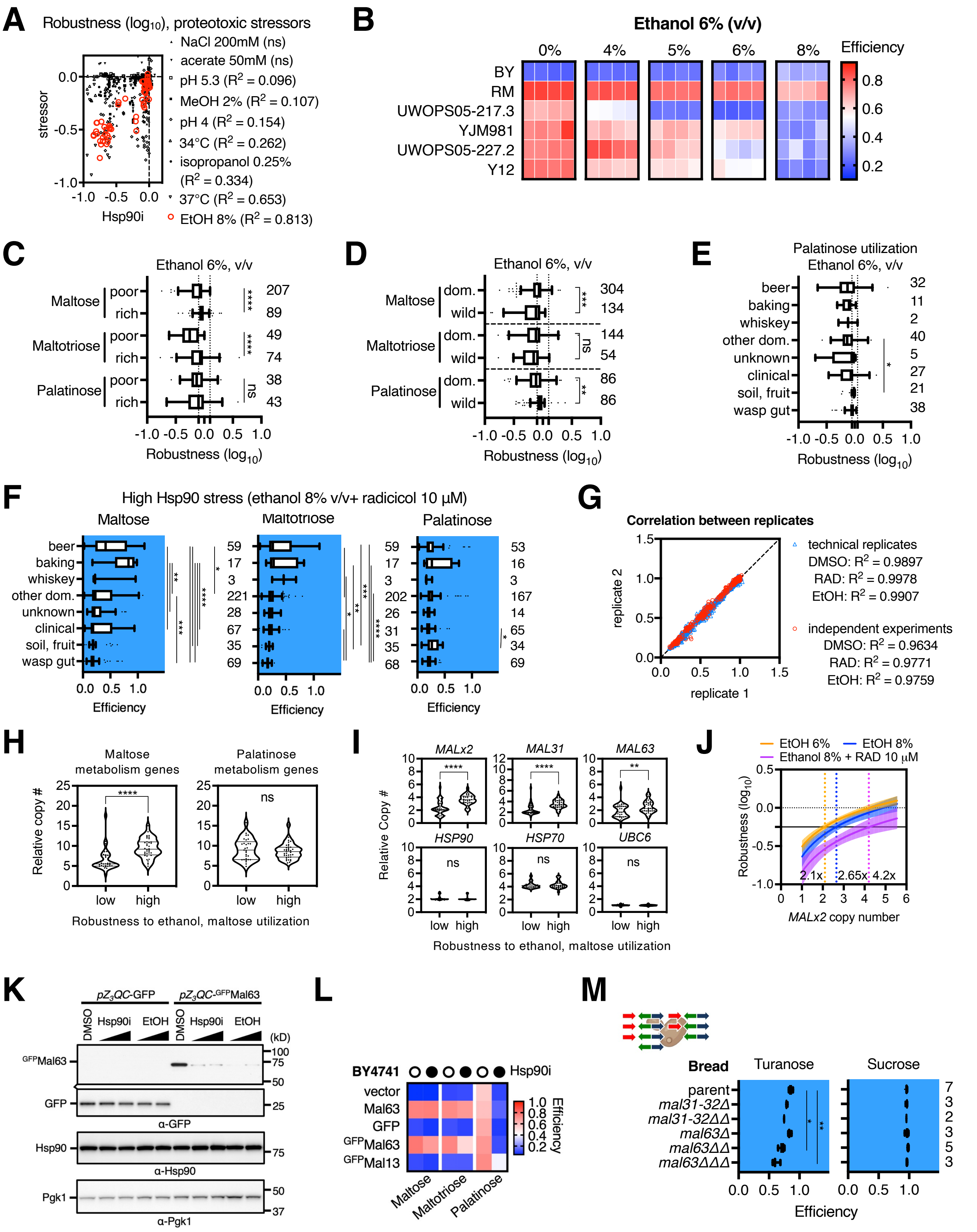

**Figure S4. Niche-related proteotoxic exposures phenocopy low-level Hsp90 inhibition in perturbing traits.** (**A**) Correlation between the effect of moderate-level Hsp90 inhibition (radicicol, 10 μM) and various stressors associated with industrial beer fermentations on maltose metabolism. "ns" indicates not significant comparisons. All other correlations were significant as per Pearson (cutoff, $p < 0.05$; 37°C, $R^2 = 0.6525$, $p < 0.0001$; ethanol 8%, $R^2 = 0.8125$, $p < 0.0001$). (**B**) Heat map of maltose utilization efficiency for 6 wild-derived *S. cerevisiae* strains under basal conditions and under increasing concentrations of ethanol (4 – 8%, v/v, compared to Hsp90i, Figure S1H). (**C-E**) Box plots of robustness of the indicated metabolic traits to moderate ethanol stress (6%, v/v) parsed by (C) maltose prevalence in domestication niche, (D) domestication (wine, beer, bread, distillation, sake, cider, etc.) *versus* wild origin (clinical, soil, fruit, wasp gut), (E) niche of strain origin. (**F**) Box plots of efficiency for the indicated metabolic traits under strong Hsp90 stress (ethanol 8% + radicicol, 10μM). Significance between distributions was determined using the Kruskal-Wallis test with Dunn's multiple comparisons. (**G**) Correlation between technical and biological replicates of relative growth under ethanol exposure (EtOH, 6 – 8%, v/v), Hsp90 inhibition (radicicol, 4 – 10 μM), and control (DMSO, low *vs*. high concentration). (**H** and **I**) Relationship between robustness to metabolic traits to (H) moderate and high (I) ethanol stress (6% *vs.* 8%, v/v) and copy number of *MAL* genes required for maltose utilization (*MAL31*, *MALx2*, *MAL63*) and palatinose utilization (*MPH2-3*, *IMA1-5*, *MAL13*, *MAL33*, *YFL052W*), excluding *MAL11* which belongs to both pathways, *versus* control genes. Significance between distributions was determined using a two-tailed Mann-Whitney test. (**J**) Interpolation plots showing threshold effect of *MALx2* relative copy number on the robustness of maltose metabolism against increasing ethanol concentrations and Hsp90 stress. For robustness values of -0.25, the 95% confidence intervals from the linear regression analysis of *MALx2* relative copy number, are ethanol 6%: 1.843 to 2.506, $R^2 = 0.6305$, $p < 0.0001$, ethanol 8%: 2.487 to 3.237, $R^2 = 0.4281$, $p < 0.0001$, ethanol 8% + Radicicol 10 μM: 3.587 to 5.002, $R^2 = 0.2909$, N = 65). (**K**) Western blotting against GFP tagged Mal63 protein expressed under the control of a β-estradiol-inducible promoter (pZ3QC) identifies changes in steady state levels upon Hsp90 stress. Log phase cultures growing in YPM supplemented with β-estradiol (100 nM) at 30°C were treated for 2 hours with Hsp90 inhibition (radicicol, 4 μM) and ethanol stress (6%, v/v) before collection and total protein precipitation. Pgk1 was used as a loading control. (**L**) Heat map of metabolic trait efficiency under basal conditions (vehicle control, open spheres) and Hsp90 inhibition (radicicol, 10 μM, closed spheres) in BY4741 strains overexpressing the indicated GFP-tagged proteins from episomal constructs. Mal13-GFP was used as control. Cells harboring episomal plasmids were grow in SD media lacking uracil. (**M**) Genotype schematic and box plots of basal efficiency of turanose and sucrose metabolism for the indicated derivatives of industrial bread strain. Values and standard deviations were derived from 2 – 5 independent biological replicates and 2 independent experiments.

**Fig. S5. Rapid selection for redundancy under ecological conditions of niche-related Hsp90 stress.**

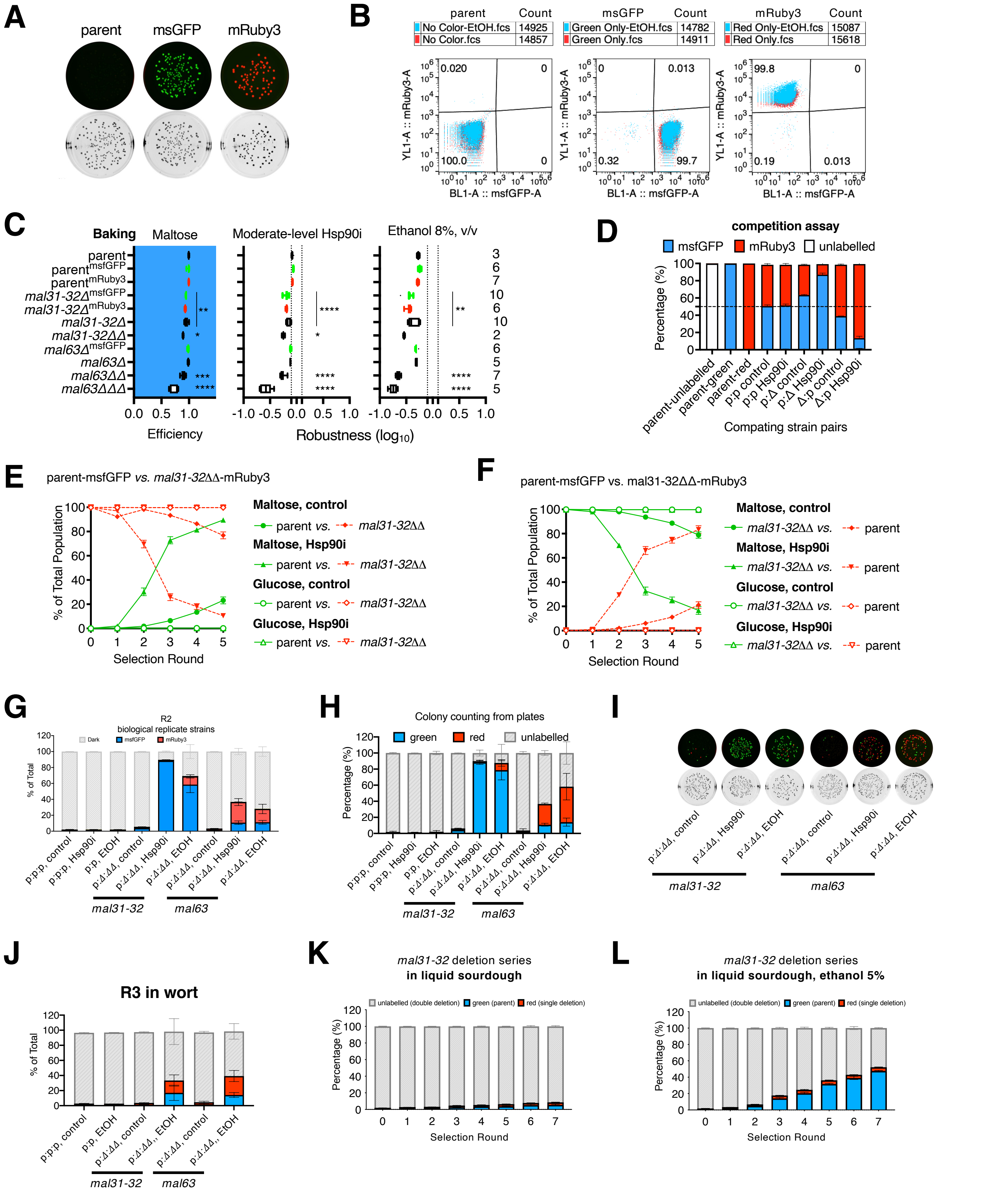

**Figure S5. Rapid selection for redundancy under ecological conditions of niche-related Hsp90 stress.** (**A**) Image of colonies of commercial bread strain and derivatives stabling expressing the indicated fluorescent proteins observed under a CCD camera after growth on SM agar plates for 3 days at 30°C. (**B**) Cell cytometry of cultures of parental and labelled derivative strains grown to saturation in liquid YPM media (red dots) *versus* YPM containing ethanol (8%, v/v, blue dots) over 3 days at 25°C. (**C**) Box plot of basal efficiency and robustness of maltose metabolism to moderate-level Hsp90 inhibition (radicicol, 10 μM) and ethanol stress (8%, v/v). (**D**) Competition assay between indicated parent and single copy *mal31-32* disrupted derivative strains competing in 1:1 mixture (green:red) in YPM under basal and low-level Hsp90 inhibition (radicicol, 4 μM) conditions. Strains were competed in both fluorescent marker combinations as indicated. (**E** and **F**) Competition assays involving 1:1,000 mixtures of the indicated strains competing over sequential bottlenecks (1:300 dilution) in (E) YPD and (F) YPM media under basal and low-level Hsp90 inhibition (radicicol, 4 μM) conditions in both marker orientations. (**G-I**) Representative biological replicate experiment of Figure 5 competition assays using independent, fully sequenced strains. (G) Cell cytometry result and (H) on plate colony counting results of selection round 2 (~10 – 13 generations). (I) Fluorescence images of representative agar plates from round 2 of this experiment. (**J**) Three-strain batch competition assay in wort and effects of ethanol exposure (8%, v/v). Batch culture 3 (R3) is shown for *mal31-32* and *mal63* disruption lineages. (**K** and **L**) Frequency plots of genotypes competing in 1:1:98 mixed cultures in liquid sourdough media under low ethanol stress (5%, v/v). Change in genotype frequencies over time (replication round) is depicted for a bread yeast-derived *mal31-32* disruption lineage completing under (K) basal conditions, and (L) low ethanol stress. Values and standard deviations were derived from 4 independent technical replicates. Values and standard deviations were derived from 2 independent replicate experiments. Significance between distributions was determined using the Kruskal-Wallis test with Dunn's multiple comparisons.
